# A conditional strategy for cell-type specific labeling of endogenous excitatory synapses in *Drosophila* reveals subsynaptic architecture

**DOI:** 10.1101/2022.10.17.510548

**Authors:** Michael J. Parisi, Michael A. Aimino, Timothy J. Mosca

## Abstract

Chemical neurotransmission occurs at specialized contacts where presynaptic neurotransmitter release machinery apposes clusters of postsynaptic neurotransmitter receptors and signaling molecules. A complex program underlies recruitment of pre- and postsynaptic proteins to sites of neuronal connection and enables the correct three-dimensional synaptic organization that underlies circuit processing and computation. To better study the developmental events of synaptogenesis in individual neurons, we need cell-type specific strategies to visualize the individual proteins at their endogenous levels at synapses. Though such strategies exist for a variety of presynaptic proteins, postsynaptic proteins remain less studied due to a paucity of reagents that allow visualization of endogenous individual postsynapses in a cell-type specific manner. To study excitatory postsynapses, we engineered *dlg1[4K]*, a conditional, epitope-tagged marker of the excitatory postsynaptic density in *Drosophila*. In combination with binary expression systems, *dlg1[4K]* effectively labels postsynaptic regions at both peripheral neuromuscular and central synapses in larvae and adults. Using *dlg1[4K]*, we find distinct rules govern the postsynaptic organization of different adult neuron classes, that multiple binary expression systems can concurrently label pre- and postsynaptic regions of synapses in a cell-type-specific manner, and for the first time, visualize neuronal DLG1 at the neuromuscular junction. These results validate a novel strategy for conditional postsynaptic labeling without the caveats of overexpression and demonstrate new principles of subsynaptic organization. The use of *dlg1[4K]* marks a notable advancement in studying cell-type specific synaptic organization in *Drosophila* and the first example of a general postsynaptic marker to complement existing presynaptic strategies.

## INTRODUCTION

Synaptic connections are highly specialized structures that mediate neurotransmission in the brain, are tightly regulated developmentally, and must maintain cohesion over the organism’s lifespan while adjusting strength and plasticity in response to environmental stimuli. Synapses underlie the control of learning, memory, cognition, and motor behavior by transmitting electrical and chemical signals within and between circuits (Mayford et al., 2012). In chemical neurotransmission, the synapse is an asymmetrical structure: the presynaptic terminal localizes the machinery needed to produce and release neurotransmitters (NT) into the synaptic cleft while the postsynaptic terminal enables neurotransmitter reception to propagate neuronal signals. After formation, synapses can undergo extensive remodeling in an activity-dependent manner, responding to experience by altering levels and subunit ratios of NT receptors, fundamentally adjusting the properties of the postsynapse. As such, understanding how synapses develop, what components comprise a mature connection, and how synaptic connections are ordered with respect to the three-dimensionality of circuits remain fundamental questions in neuroscience. Further, a better understanding of synaptic development can inform our understanding of what is perturbed in neurodevelopmental and neurodegenerative disease conditions (Henstridge et al., 2016).

To understand how synapses function and form, we must distinguish, image, and target both the pre- and postsynaptic regions during different developmental stages, with cell-type specificity. The dynamic nature of synaptic biology requires techniques to visualize and manipulate these processes. Historically, peripheral synapses like the neuromuscular junction (Harris and Littleton, 2015; Sanes and Lichtman, 1999) have been powerful systems for studying synaptic cell biology as these synapses are large, experimentally accessible, and as a result of their size and organization, have readily separable pre- and postsynaptic terminals. This allows the use of antibodies to study synaptic biology with the caveat that they cannot distinguish between proteins present in specific neurons or between pools of that protein at the pre- or the postsynapse. In the central nervous system, however, the added density of synaptic connections, the increase in the diversity of neuronal types, and the proximity of pre- and postsynaptic terminals, precludes the use of antibodies to perform high-resolution cell-type specific analyses *in vivo* (Duhart and Mosca, 2022). To date, though, a considerable body of ultrastructural, biochemical, and proteomic studies identified many of the principle protein components that make up the vertebrate and invertebrate synapse (Helm et al., 2021; Wilhelm et al., 2014). These studies enabled the creation of a host of genetically encoded reagents aimed at labeling specific synaptic proteins and, when combined with binary expression systems or viral-mediated delivery, enable cell-type specificity for studying circuits (Duhart and Mosca, 2022). The presynaptic proteome especially has validated many previously identified components of the active zone, a site specialized for neurotransmitter release, and added to a growing set of useful conserved immunohistochemical reagents (Boyken et al., 2013; Gronborg et al., 2010; Morciano et al., 2005, 2009; Weingarten et al., 2014). In *Drosophila*, the protein Bruchpilot (Wagh et al., 2006), a highly conserved ortholog of the vertebrate CAST protein (Ohtsuka et al., 2002), is widely used as a presynaptic marker immunohistochemically and via various genetically encoded constructs to study presynapses in distinct neuronal populations with cell-type specificity as these constructs can label the endogenous active zone and do not interfere with its function (Chen et al., 2014; Christiansen et al., 2011; Coates et al., 2017; Duhart and Mosca, 2022; Fouquet et al., 2009; Kremer et al., 2010; Mosca and Luo, 2014; Mosca et al., 2017; Urwyler et al., 2015; Wagh et al., 2006). Studies using Bruchpilot are routinely complemented with additional cell-type specific strategies to visualize synaptic vesicles, trafficking proteins, and other presynaptic components (Certel et al., 2022a, 2022b; Duhart and Mosca, 2022; Venken et al., 2011).

In contrast, fewer strategies exist to generally label endogenous postsynaptic sites in a cell-type specific manner without the caveats associated with overexpression due to unique challenges inherent to the postsynapse. Excitatory and inhibitory neurotransmission utilize the same presynaptic release site machinery while employing different vesicular transporters specific to the neurotransmitter in question (for example, glutamate for excitatory neurotransmission and GABA for inhibitory neurotransmission). Therefore, presynaptic labeling strategies that visualize release site machinery will be able to capture both excitatory and inhibitory synapses. Postsynaptic labeling is more challenging as receptors for excitatory and inhibitory neurotransmission are not shared. Indeed, each neurotransmitter has a fundamentally distinct receptor so even within one class of neurotransmission (excitatory vs. inhibitory), strategies that label one receptor would necessarily omit others. There is a strong need for general cell-type specific labels of excitatory and inhibitory postsynaptic terminals. Further, such labels should be cell-type specific to label postsynaptic terminals in a select class of cells, circumventing the issue of increased density in the central nervous system. In excitatory neurons, the postsynapse is marked by an electron dense structure called the postsynaptic density (PSD) positioned adjacent to presynaptic active zones (Gray, 1959). The vertebrate PSD has been extensively characterized (Bayés et al., 2011, 2014; Biesemann et al., 2014; Collins et al., 2006; Gray, 1959) and is densely packed with neurotransmitter (NT) receptors, signaling molecules, and PSD-95, an abundant scaffolding protein (Cho et al., 1992). PSD-95 interacts with several key PSD components and serves as a central scaffold to organize NT receptors (Won et al., 2017). In *Drosophila*, the PSD-95 orthologue *discs large1* (*dlg1*) (Woods and Bryant, 1991) is well-studied at peripheral synapses as a label of excitatory postsynaptic neuromuscular junctions (NMJs) (Budnik et al., 1996; Harris and Littleton, 2015) with a role in synaptic maturation, NT organization, and neurotransmission. DLG1 / PSD95 proteins are highly conserved and extensively used as immunohistochemical markers to study synaptic development and function. In *Drosophila*, DLG1 labeling has been crucial (Thomas et al., 2000) for nervous system study but antibody-based methods have the limitation of lacking the specificity required to examine synaptic connections in a select group of cells or to map a specific part of a neuronal circuit. Strategies for general cell-type specific postsynaptic labeling, however, have lagged behind those for presynaptic labeling. DLG1 / PSD-95 overexpression alters basal synaptic development, structure, and function in both *Drosophila* and mammalian systems (Ehrlich and Malinow, 2004; Ehrlich et al., 2007; Elias et al., 2008; Gorczyca et al., 2007; Sturgill et al., 2009). Although this approach has yielded considerable information about PSD-95 function and binding partners, it precludes the utility of overexpression for examining basal synaptic function and development. An optimal labeling strategy would allow DLG1 / PSD-95 to be expressed at endogenous levels and still be labeled in a cell-type specific manner. In recent years, a number of cell-type specific approaches allowing the study of postsynaptic proteins using endogenous levels based on technologies like intrabody binding (Gross et al., 2013) and CRISPR / Cas9 modification (Fang et al., 2021; Nishiyama et al., 2017; Willems et al., 2020) enabled cell-type specific labeling of many synaptic proteins, including postsynaptic proteins like PSD-95, allowing unprecedented study – the advantage of the sparse labeling nature of such techniques, however, is also a disadvantage, as it does not allow for labeling of all the proteins contributed by a class of neurons. A strategy is also needed to label postsynaptic proteins in all neurons of a single class, at their endogenous levels, to assess contributions of all cells of interest to a circuit rather than a portion of those cells. Conditional labeling in *Drosophila* fills some of this need using recombineering (Chen et al., 2014), FlpTag to conditionally label specific proteins (Fendl et al., 2020) in cells where exogenous FLP recombinase is provided, or reconstituted GFP labeling (Feinberg et al., 2008; Kamiyama et al., 2021; Kondo et al., 2020) using the split GFP system to label CRISPR-modified genes. To date, though, such strategies have labeled select postsynaptic receptors and not general postsynapses, limiting the broad applicability of these tools.

To develop a cell-type specific endogenous label that generally encompasses excitatory postsynapses, we designed a CRISPR-mediated modification of the *dlg1* locus in *Drosophila*. This novel strategy uses FLP recombination to introduce an epitope tag at the C-terminus of the DLG1 protein while maintaining endogenous protein levels. Binary expression systems like GAL4, lexA, or QF conditionally provide FLP recombinase to label endogenous DLG in a cell-type-specific manner. We applied this new approach, *dlg1[4K]*, to examine synaptic architecture at the peripheral NMJ and at central neuron synapses in the olfactory system. We demonstrate robust epitope signal at multiple classes of synapses, utility with the major binary expression systems in *Drosophila*, and strategies for simultaneous pre- and postsynaptic labeling using multiple expression systems concurrently. Finally, we use *dlg1[4K]*, to reveal previously unappreciated subsynaptic architecture of DLG1 at central and peripheral synapses. We anticipate that this strategy will be applicable at all excitatory *Drosophila* synapses and provide an additional, generalized strategy for cell-type specific labeling of synaptic proteins.

## RESULTS

### Rationale and design of a knock-in at the dlg1 locus

The absence of a general postsynaptic label that can be utilized in a cell-type specific fashion has stymied progress in studying synaptic organization in *Drosophila*. To overcome the lack of such a marker, we targeted the *dlg1* locus, which encodes the fly homologue of PSD-95, a postsynaptic scaffolding molecule in most classes of excitatory neurons (Budnik et al., 1996; El-Husseini et al., 2000; Woods and Bryant, 1991). *Drosophila* DLG1 belongs to the MAGUK (<u>m</u>atrix <u>a</u>ssociated <u>gu</u>anylate <u>k</u>inase) family of proteins and is encoded by a genomic region that spans ~40 kB and produces 21 annotated transcript isoforms using alternative promoters and start codons that can produce non-overlapping proteins (Graveley et al., 2011; Mendoza et al., 2003) (Fig. S1). The largest DLG1 isoform includes four protein-protein interaction motifs (L27, PDZ, SH3 and GK) that mediate a diverse array of protein-protein interactions (Doerks et al., 2000; Feng et al., 2005; Petrosky et al., 2005; Sheng and Kim, 2011). The single fly *dlg1* gene is highly conserved across species with five *dlg1*-related genes in humans. DLG1 plays essential roles in neuronal function (Budnik et al., 1996; Mendoza et al., 2003) as well as planar cell polarity in epithelial tissues, cell-to-cell adhesion as a principle component of the septate junction complex (Abbott and Natzle, 1992; Woods and Bryant, 1991; Woods et al., 1996), and is hypothesized to have a role in vesicle function as well (Walch, 2013).

We designed a strategy that would conditionally label DLG1 only in cells of interest at endogenous levels to avoid any artifacts from DLG1 overexpression (Fig. 1). To do this, we took advantage of the FLP / FRT recombinase system (Dang and Perrimon, 1992) and specifically, the FRT-STOP-FRT cassette (Weasner et al., 2017). This strategy places a transcriptional stop sequence between two FRT recombination sites. In the absence of a FLP recombinase, the ribosome will read through the first FRT site and halt transcription at the stop sequence (Golic and Lindquist, 1989; Xu and Rubin, 1993). In the presence of FLP recombinase, though, site-specific recombination will occur between the FRT sites, removing the stop sequence, and allowing for continued readthrough of the open reading frame. We placed a V5 epitope tag immediately following the FRT-STOP-FRT sequence (FRT-STOP-FRT-V5) so that in the presence of FLP, a V5 tag would be expressed (Fig. 1B). We reasoned that, based on the exonic structure of *dlg1*, that inserting this cassette immediately upstream of the most 3’ stop codon would result in a modified *dlg1* ORF (Fig. 1C): in the absence of FLP, the ORF would encode an untagged, full-length protein (Fig. 1D) and in the presence of FLP, would encode a V5-tagged full-length protein (Fig. 1E), both under the control of their endogenous promoter (Fig. 1). Thus, the cell-type specificity of DLG1 labeling could be achieved by providing FLP only in select cells; a selected driver line (via GAL4, lexA, QF) expressing FLP would remove the stop cassette resulting in DLG1 protein translated in frame with a V5-tag expressed under the control of the endogenous promoter only in those cells expressing FLP (Fig. 1E).

**Fig. 1.**
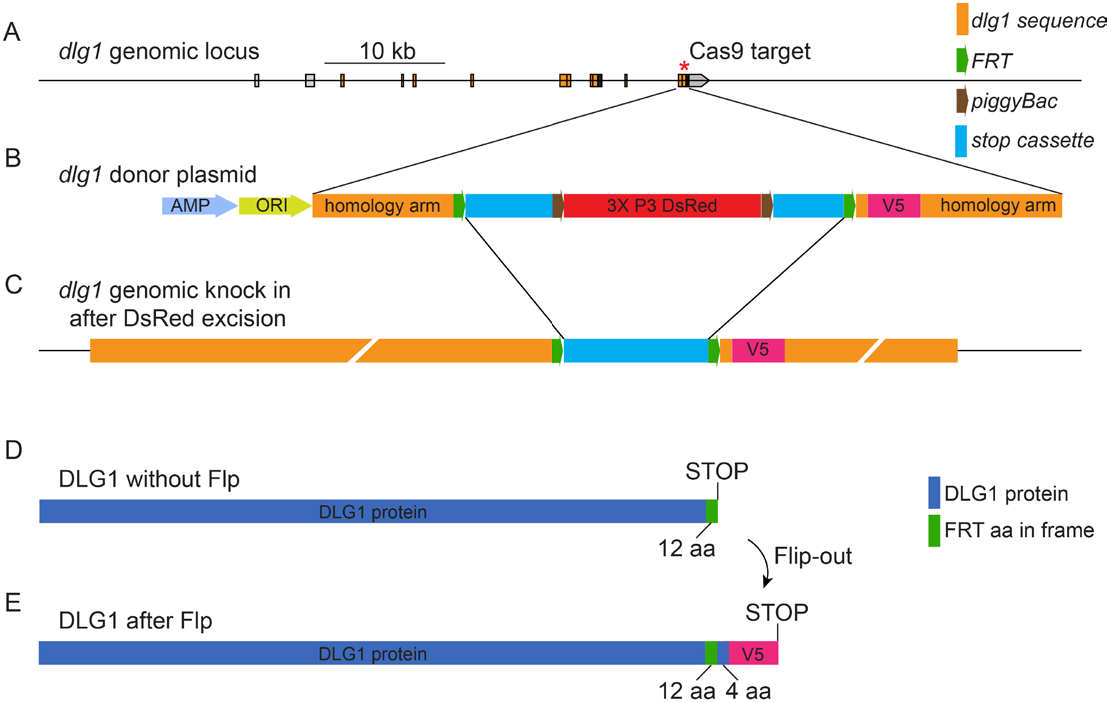
Strategy for engineering a conditionally inducible V5 epitope tag at the *dlg1* locus. (A) Map of the *dlg1* genomic locus with transcript dlg1-RB depicted. Integration was directed to the distal-most stop codon (indicated by an asterisk). (B) The donor construct was built into a plasmid backbone with two 500 bp *dlg1* homology arms flanking the FRT-STOP-FRT cassette and a V5 epitope tag. A DsRed cassette flanked by piggyback transposon inverted repeats (brown arrows) is inserted within the FRT-STOP-FRT sequence. (C) Representation of the *dlg1* locus knock-in after scarless excision of the 3X P3 DsRed cassette. (D) Schematic of the translated product of the dlg1 locus in the absence of a FLP recombinase. Translation of DLG1 under the control of the endogenous promoter but without FLP present produces a DLG1 product with 12 additional amino acids (due to the FRT site) but lacking the last 4 DLG1 amino acids and the V5 epitope tag. A UAA stop codon was engineered directly 3’ of the FRT sequence. (E) Schematic of the translated product after a FLP-out event shows DLG1 reading in frame through the FRT sequence and into the V5 tag. This results in a product with 54 amino acids added to DLG1 (due to the FRT site) and the V5 epitope tag.

To make *dlg1-FRT-STOP-FRT-V5*, we used CRISPR / Cas9 genomic engineering with Homology Directed Repair (Bier et al., 2018; Gratz et al., 2013) and edited the *dlg1* locus with a conditional FLP-mediated epitope tag at the most distal stop codon (Fig. S1). The construct design uses a FLP-out strategy (Golic and Lindquist, 1989; Xu and Rubin, 1993) with a transcriptional stop cassette flanked by tandem minimal FRT sites (Nern et al., 2015) that we modified by adding the sequence encoding a V5-epitope immediately following the FRT-STOP-FRT cassette and then finally by adding two flanking 500 bp homology arms encoding *dlg1* sequence at the 5’ and 3‘ ends (Fig. 1B). A single nucleotide (C) was added directly between the left homology arm and the 5’ FRT to maintain the reading frame; in the absence of FLP, 12 residues (from the FRT sequence) are added to the DLG1 protein (Fig. 1D). A UAA stop codon was also engineered directly after the FRT sequence. We further inserted a 3xP3-dsRed marker flanked by piggyBac transposon sites within the transcriptional stop sequence to provide a marker that could be scored visually (red fluorescence in the eye) (Fig 1B). In the presence of a FLP-out event, 12 amino acids from the FRT site plus 4 additional DLG1 residues between the FRT site and the V5 epitope are included in the ORF (Fig. 1E). Following the V5 epitope, the endogenous stop codon and 3’ UTR for DLG1 are retained, ensuring that cessation of the open reading frame via its native stop sequence.

We used CRISPR / Cas9 engineering with dlg1-specific gRNAs via germline transformation to introduce this construct into the *Drosophila* genome (Gratz et al., 2013, 2015; Port et al., 2014). The resultant flies were then crossed to a piggyBac transposase source (Horn et al., 2003) to remove the 3xP3-dsRed label in a “scarless” fashion, resulting in the final FRT-STOP-FRT-V5 construct within the *dlg1* locus. This line was termed *dlg1[4K]* and used for all subsequent experiments. Flies bearing the inserted sequence with the 3xP3-dsRed removed were homozygous viable and fertile, suggesting that this manipulation of the endogenous dlg1 locus did not preclude the essential function of the gene (Woods and Bryant, 1991). Moreover, in all experiments, FLP events with *dlg1[4K]* did not influence viability, nervous system structure, developmental timing, or adult behavior. This indicates that the manipulations do not interfere with endogenous DLG1 function. Thus, the *dlg1* locus is amenable to genomic manipulation to produce a conditional, FLP-inducible epitope tag for cell-type specific labeling.

### Muscle induced dlg1[4K] labels the NMJ and co-localizes with endogenous Dlg1

To test cell-type specific labeling using *dlg1[4K]*, we first turned to the larval neuromuscular junction (NMJ), a widely used and readily tractable model for studying synaptic development and architecture (Keshishian et al., 1996). Each segment of the third instar larva contains a repeated pattern of body wall muscles that are innervated by motor neurons with stereotyped projections of strings of synaptic boutons (Hoang and Chiba, 2001). DLG1 is notably expressed at the NMJ where it plays many diverse roles in postsynaptic organization, synaptic maturation, and development (Astorga et al., 2016; Budnik et al., 1996; Mendoza et al., 2003; Parnas et al., 2001; Thomas et al., 2000). DLG1 labels the region of the bouton immediately surrounding the presynaptic membrane but does not extend completely into the cytoskeletal shell surrounding the bouton (Budnik et al., 1996; Lahey et al., 1994; Pielage et al., 2006; Restrepo et al., 2022; Wang et al., 2011). Though commonly used as a postsynaptic marker, there is genetic evidence that DLG1 can also function presynaptically at the NMJ (Astorga et al., 2016; Budnik et al., 1996; Mendoza et al., 2003) though the majority of its functions are postsynaptic. Antisera against DLG1 are thus commonly used to study synaptic development at the NMJ. Further, due to the structure of the NMJ, there is ready separation between the pre- and postsynaptic compartments allowing for high-resolution imaging. Therefore, we reasoned that inducing *dlg1[4K]* labeling throughout the entire larval musculature should recapitulate DLG1 immunostaining at the NMJ. To do so, we first utilized GAL4, QF, or lexA driven via the *Mef2* (*Mef2-GAL4*, *Mef2-lexA*, or *Mef2-QF2*) promoter (Lilly et al., 1995) to express the FLP recombinase (via UAS-FLP, lexA-op-FLP, or QUAS-FLP) in all muscles. When we did this in the background of the *dlg1[4K]* insertion, this removed the FRT-STOP-FRT cassette only in muscles where the FLP was provided, enabling DLG1 to be labeled with the V5 tag. We stained the resultant larvae with antibodies to V5 and to the endogenous DLG1 to label postsynaptic DLG1 and with antibodies to HRP (Jan and Jan, 1982) to label the presynaptic motoneuron. In non-expressing controls (containing a FLP transgene and the *dlg1[4K]* insertion), we observed robust labeling with DLG1 and HRP, as expected, but failed to observe any V5 staining (Fig. 2A, C, E). However, when FLP was expressed in postsynaptic muscles using the respective drivers, we observed robust V5 staining that colocalized precisely with the DLG1 staining, suggesting that the same pool of DLG was recognized by both V5 and DLG1 antisera (Fig. 2B, D, F). This indicates that epitope tag labeling of *dlg1[4K]* recapitulates the endogenous staining pattern providing proof-of-principle of our technique. Further, this indicates that expression of the knock-in construct is specific to cells bearing FLP expression (here supplied by a binary expression system driver). We validated these results with an independent GAL4 driver that also expresses in all muscles (*24B-Gal4*, Fig. S2A).

**Fig. 2.**
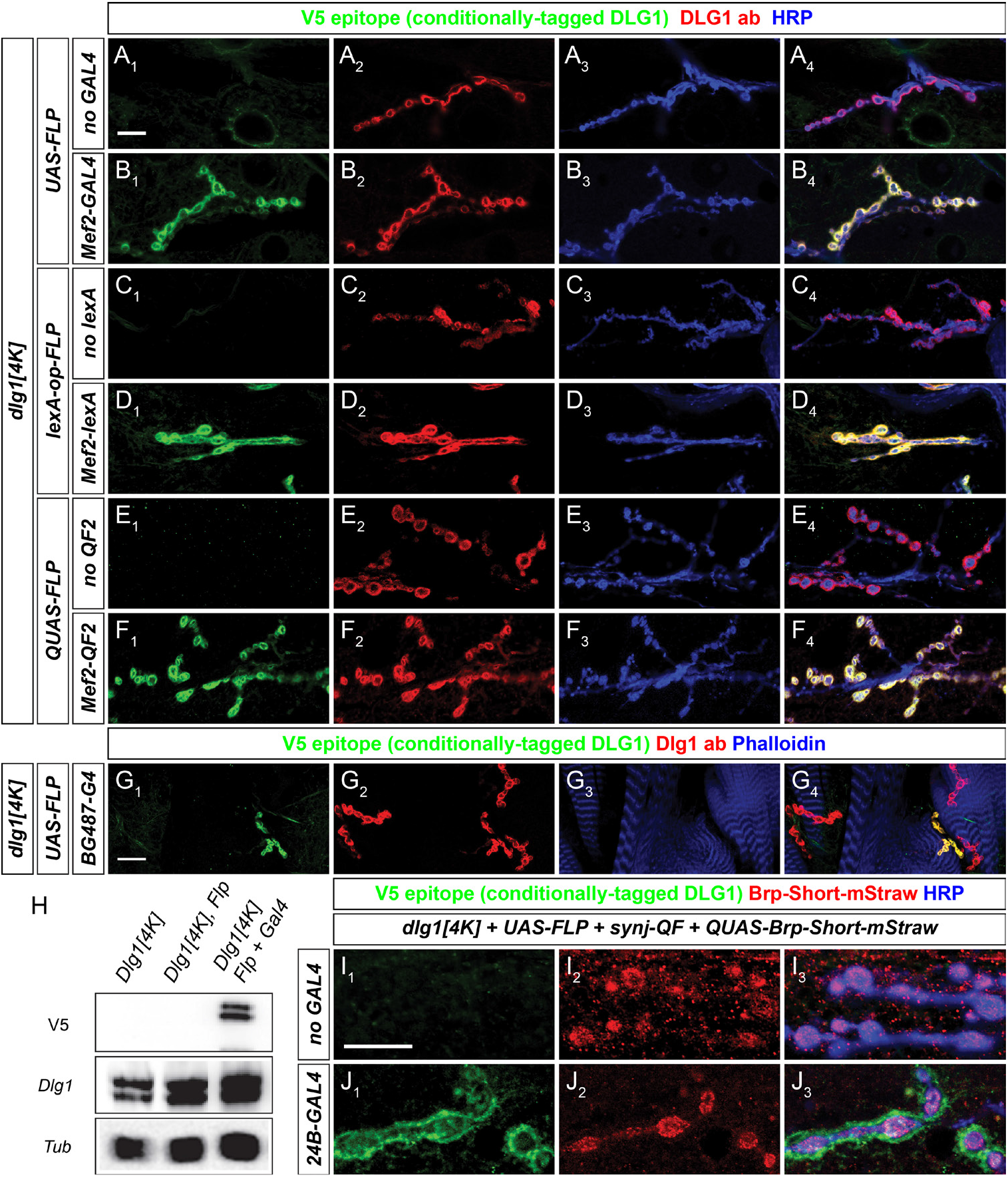
Muscle-specific labeling of DLG1 using *dlg1[4K]* via tissue-specific FLP expression. (A–G) Representative confocal images of individual NMJs in multiple genotypes and stained with antibodies to the V5 epitope (green), endogenous DLG1 (red), and the neuronal marker HRP (blue). Negative controls lacking the expression driver (A, no GAL4; C, no lexA; E, no QF2) but containing the dlg1[4K] allele and a FLP transgene show no V5 immunoreactivity while endogenous DLG1 is detected. When the respective binary expression is present and driven by a muscle-specific promoter, *Mef2* (B, *Mef2-GAL4*; D, *Mef2-lexA*; F, *Mef2-QF2*), V5 immunoreactivity is present at the NMJ that precisely overlaps with endogenous DLG1 staining. This indicates that endogenous muscle DLG1 is now labeled with a V5 tag, due to the presence of *dlg1[4K]* and FLP recombinase. (G) Representative confocal images of *dlg1[4K]* larvae expressing FLP in only a subset of larval muscles and stained with antibodies to DLG1-V5 (green) and endogenous DLG (red) with phalloidin (blue). In select terminals that coincide with GAL4 expression, DLG1-V5 expression is evident while in others, only endogenous DLG1 signal is observed, demonstrating that DLG1-V5 expression is tightly linked to GAL4-driven FLP expression. (H) Western blot analysis of larval lysates from multiple genotypes demonstrating immunoreactivity of endogenous DLG1 and the V5 epitope. Endogenous DLG1 is visible in all genotypes but V5 is only visible when *dlg1[4K]* and FLP are present, suggesting no leaky expression. Tubulin is used as a loading control. (I-J) Representative confocal images of multiple genotypes stained with antibodies to DLG1-V5 (green), dsRed (red, to recognize Brp-Short-mStraw), and HRP (blue). Multiple cell types can be labeled in the same sample via the use of multiple binary expression systems. Pan-neuronal QF2 labels motoneurons while DLG1-V5 is only present in the muscle when both a muscle GAL4 source and UAS-FLP are combined. Scale bars, 10 μm.

Since expression of *dlg1[4K]* showed precise co-localization with endogenous DLG1 when FLP was supplied in all muscles, we next sought to separate the two signals within the same animal using drivers where GAL4 expression is restricted to a subset of muscles. We used the *BG487-GAL4* line to express FLP in a restricted subset of larval NMJs in the anterior-most segments (Budnik et al., 1996). We observed colocalization of the V5 epitope and endogenous DLG1 only in those muscles expressing FLP while adjacent NMJs that lacked expression of FLP were only recognized by antisera to the endogenous DLG1 (Fig. 2G). Thus, *dlg1[4K]* successfully labels DLG1 only in the precise cellular pattern where FLP is supplied (here by a binary expression driver).

We also sought to determine whether any ‘leaky’ expression of the V5 epitope could be observed in the absence of FLP. Using a pan-muscle GAL4 driver, we systematically eliminated the driver and/or FLP components (Fig. S3). This test for ‘leaky’ expression showed that *dlg1[4K]* alone, or *dlg1[4K]* with UAS-FLP (but no GAL4) displayed no observable α-V5 staining (Fig. S3A-B). We further validated these results using Western blot analysis of analogous larval lysates (Fig. 2H). In each case, V5 expression via Western was only observed when FLP was actively expressed in muscles and not in any GAL4- or UAS-only controls (Fig. 2H). This indicates that there is little to no leaky expression associated with *dlg1[4K]* and that labeling is tightly coupled to the presence or absence of FLP recombinase.

The above experiments validated the *dlg1[4K]* label using a single binary expression system. However, it is often advantageous to use multiple binary expression systems simultaneously to manipulate one cell-type and label a second or to label two different cell types in tandem. Therefore, we sought to determine if *dlg1[4K]* could specifically label the NMJ postsynapse while also using a second binary expression system to concurrently label a presynaptic marker. We combined the GAL4 and QF systems to label the post- and presynaptic NMJ, respectively. We used a pan-neuronal QF driver (*synj-QF*; Petersen and Stowers, 2011) to express QUAS-Brp-Short-mStraw (Mosca and Luo, 2014), a presynaptic active zone label, in motoneurons that innervate the NMJ. Simultaneously, we used the pan-muscle driver 24B-GAL4 to drive UAS-FLP expression in muscle cells along with *dlg1[4K]* to label only postsynaptic muscle DLG1 at the NMJ. In the absence of GAL4 (Fig. 2I), we failed to observe DLG1-V5 labeling, as expected though QF-driven labeling of Brp-Short was evident (Fig. 2I). However, in the presence of GAL4, there was clear DLG1-V5 labeling that was postsynaptic to the QF-driven Brp-Short (Fig. 2J). Larval NMJs showed patterns of apposition as expected from established pre- and postsynaptic localization (Fig. 2J). Importantly, the two staining patterns did not overlap, suggesting that using both binary expression systems simultaneously did not sacrifice the cell-type specific expression aspects of each system. This labeling experiment demonstrates the utility of combining binary expression systems with *dlg1[4K]* in concert with a wide range of responders to label multiple cell types simultaneously for more sophisticated experiments. Taken together, these results suggest that *dlg1[4K]* is a highly efficient, conditional label of endogenous DLG1 with minimal leak and indicates that it can be successfully used as a labeling tool for studying DLG1 *in vivo* and with multiple binary expression systems while maintaining control of DLG1 expression via its endogenous promoter.

### dlg1[4K] labels synaptic regions in the larval central nervous system

As *dlg1[4K]* works as a postsynaptic marker at the NMJ using muscle-specific drivers to express the FLP recombinase, we next sought to extend the functionality of this tool to the larval central nervous system. We specifically examined the larval ventral nerve cord (VNC) as the center of the nerve cord contains a neuropil rich region containing synapses amongst interneurons, motoneurons, and other sensory neurons. To examine *dlg1[4K]* expression in the VNC, we used pan-neuronal GAL4 (*C155-GAL4*; Lin and Goodman, 1994) or QF (*synj-QF*; Petersen and Stowers, 2011) driver lines to express FLP in all neurons via their respective UAS- or QUAS-FLP transgenes (Fig. 3A-D). In both cases, we observed robust V5 epitope tag staining when FLP was present (Fig. 3B, D) and little to no staining in controls without FLP (Fig. 3A, C) demonstrating that V5 expression was tightly linked to the presence of FLP. V5 staining was visible in the neuropil region and was largely excluded from the cortex region of the VNC, indicating that DLG1-V5 staining was evident in the synapse-rich region and not the cell body region, as expected for a synaptic marker. DLG1-V5 was also consistent with previous work showing DLG1 staining in the VNC (Budnik et al., 1996). Moreover, DLG1-V5 staining showed regional (but not precise) colocalization with presynaptic markers Brp or CSP, suggesting that the two were apposed markers, which is consistent with DLG1 acting largely as a postsynaptic label. Though we cannot rule out a presynaptic DLG1 contribution at this resolution, the staining is consistent with DLG1-V5 localization at the synapse and accurately recapitulates the endogenous staining pattern of DLG1, demonstrating the utility of *dlg1[4K]* as a conditionally-inducible label for central synapses.

**Fig. 3.**
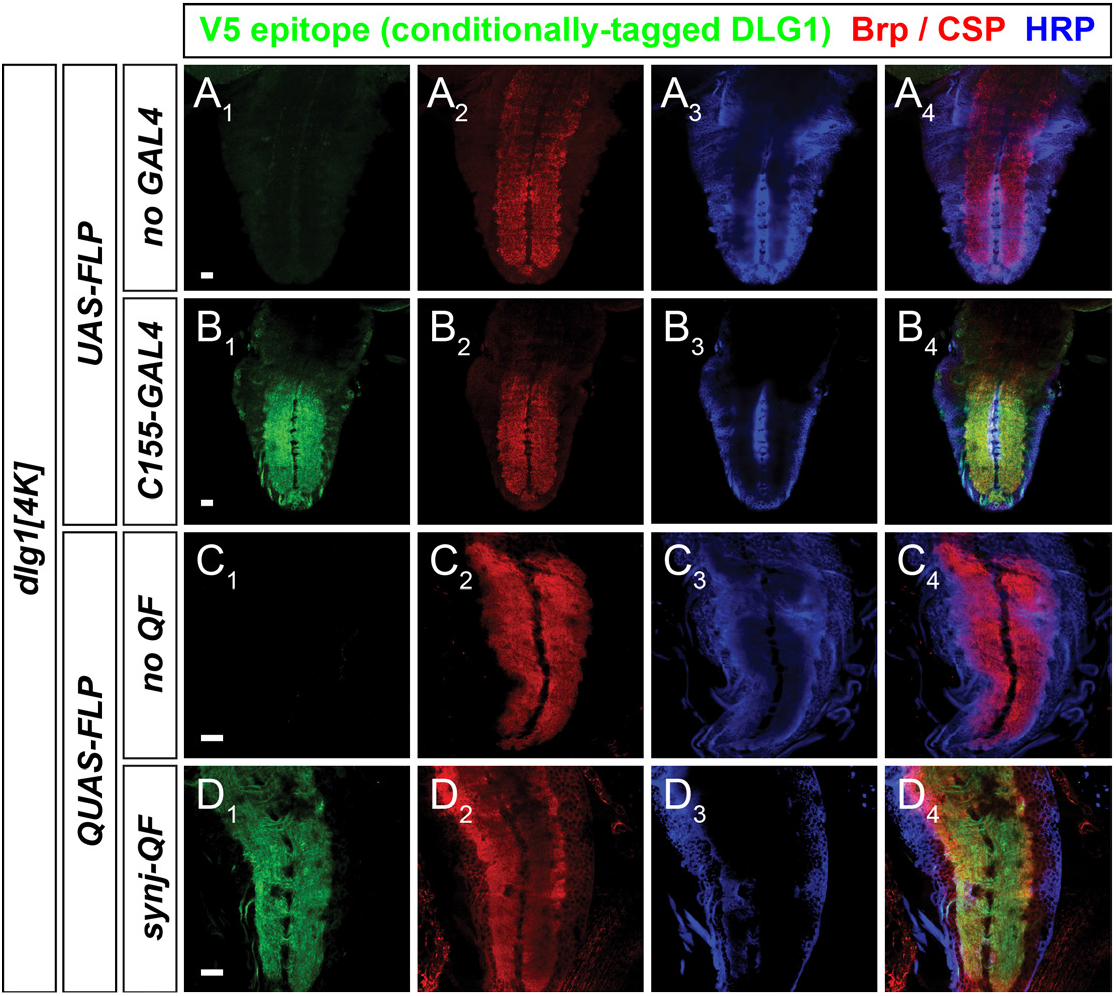
Cell-type specific labeling of DLG1 in larval neurons using *dlg1[4K]*. (A – D) Representative confocal images of larval ventral nerve cords and stained with antibodies to V5 (green, A-D), Brp (red, A-B), CSP (red, C-D), and HRP (A-D). In the absence of a GAL4 (A) or QF (C) driver, FLP cannot catalyze the recombination event in *dlg1[4K]* and no V5 labeling is observed. When the driver is present to express FLP pan-neuronally (B, D), robust V5 labeling consistent with postsynapses is observed. Scale bars, 20 μm.

### dlg1[4K] labels postsynaptic regions in olfactory neurons of the adult CNS

We next sought to examine the utility of *dlg1[4K]* in the adult *Drosophila* CNS and turned to the olfactory system and specifically, the fly antennal lobe. The fly antennal lobe is a useful model for studying wiring decisions (Jefferis and Hummel, 2006) and recently emerged as a genetic model for studying synapse development with high resolution (Duhart and Mosca, 2022; Mosca and Luo, 2014; Mosca et al., 2017). The *Drosophila* antennal lobe (AL) is a complex, yet tractable sensory circuit representing the first order processing center in the fly brain for olfactory information. Three major classes of neurons comprise the AL: olfactory receptor neurons (ORNs), projection neurons (PNs), and local interneurons (LNs). ORNs are the first order neurons whose cell bodies are housed in external structures like the antenna and the maxillary palp and project their axons to the AL. These cells are responsible for conveying olfactory information from the outside environment into the AL. There, ORNs synapse onto PNs and LNs in approximately ~50 sub-regions of each AL called glomeruli. The PNs are the second order neurons that receive signal from the ORNs and convey that to higher order olfactory processing centers in the brain like the mushroom body and the lateral horn. The LNs remain the least studied of the three neuronal classes that comprise the AL, but are widely thought to mediate gain control and inter-glomerular communication (Berck et al., 2016; Chou et al., 2010; Hong and Wilson, 2015; Liou et al., 2018; Yaksi and Wilson, 2010). Because of the tractable connectivity of these three classes of neurons, the ease of high resolution imaging, and the clear repertoire of behavioral connections to understanding synaptic biology in the AL, it remains a powerful system for understanding synaptic development, function, and organization (Duhart and Mosca, 2022). To date, the synaptic organization of the antennal lobe has largely been studied using presynaptic markers like Brp-Short (Mosca and Luo, 2014; Mosca et al., 2017) but study of postsynaptic organization has lagged behind due to a dearth of tools (Duhart and Mosca, 2022).

In the antennal lobe, a significant portion of postsynaptic terminals represent ORN axon terminals projecting onto PN dendrites. Therefore, we first tested *dlg1[4K]* using projection neurons and specifically, *GH146-GAL4*, which drives expression in 2/3 of all AL PNs. Endogenous DLG1 immunoreactivity recognizes the entire AL (Fig. 4A) but in the absence of GAL4-driven FLP, no V5 immunoreactivity is observed, consistent with the tight control of *dlg1[4K]* activity without recombination. However, when FLP is provided in PNs using *GH146-GAL4* (Berdnik et al., 2008), we observed V5 immunoreactivity that directly overlapped with most endogenous DLG1 staining (Fig. 4B), suggesting that *dlg1[4K]* successfully recapitulates DLG1 expression in PNs. Importantly, there was not complete overlap, as the DLG1 antibody recognizes all contributions of DLG1 while FLP only catalyzes recombination in *GH146*-positive PNs. Such contributions could be from other neuronal populations but may also be representative of DLG1 involved in septate junctions and/or in glia. We observed similar results when co-staining with antibodies against Brp, a presynaptic active zone marker (Fig. 4C-D). Importantly, there was little overlap between the Brp and V5 staining in GH146-positive PNs when FLP was present, rather, the two signals were closely apposed (Fig. 4D) as expected for pre- and postsynaptic markers. This again suggests that *dlg1[4K]* successfully recapitulates endogenous DLG1 expression and that the majority of DLG1 in the PNs represents postsynaptic architecture.

**Fig. 4.**
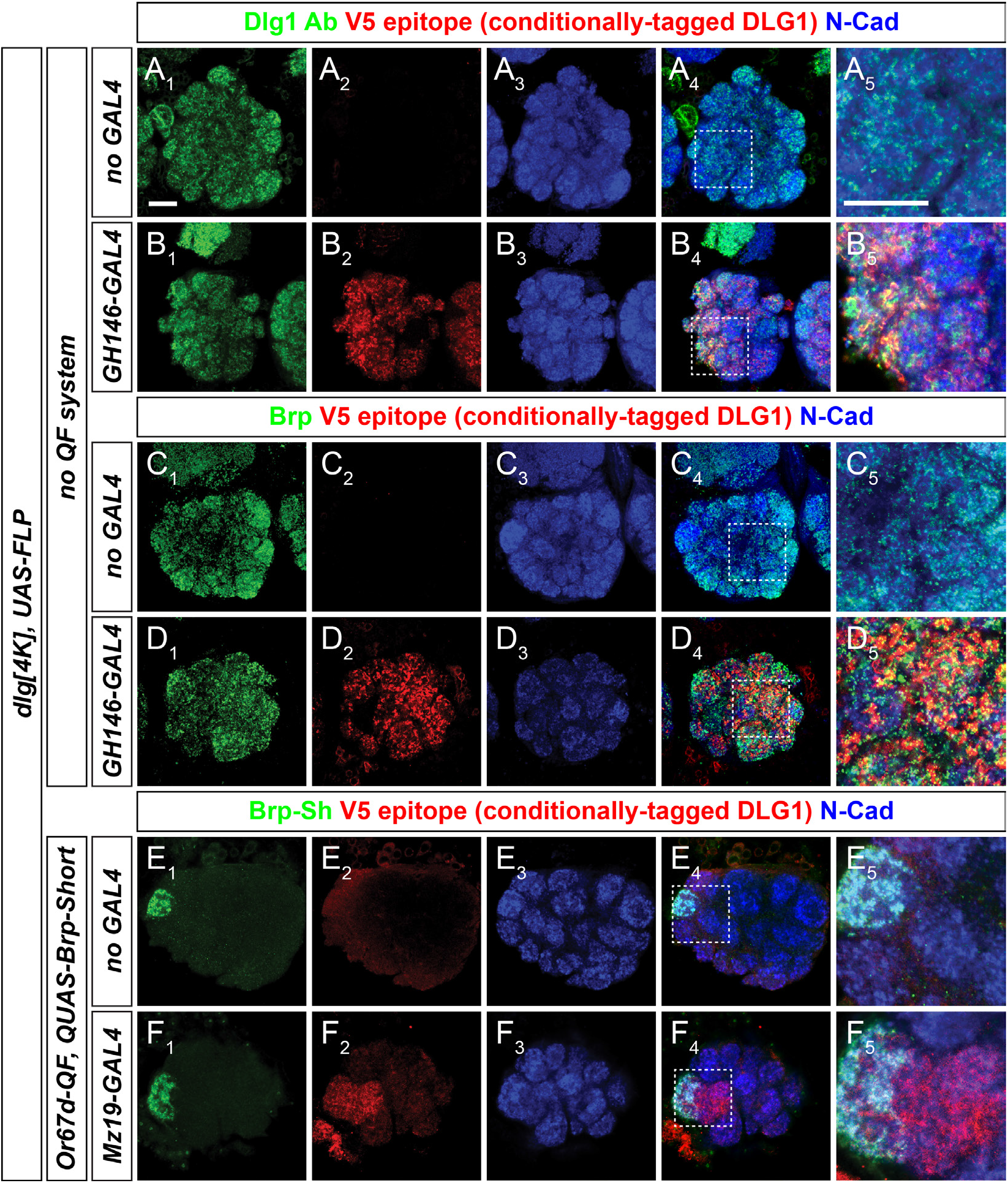
Cell-type specific FLP expression with *dlg1[4K]* labels postsynapses in adult olfactory neurons of the central nervous system. (A-B) Representative confocal images of adult antennal lobes of multiple genotypes stained with antibodies to endogenous DLG1 (green), DLG1-V5 (red), and N-Cadherin (blue). In the absence of GAL4 (A), no *dlg1[4K]* expression is observed while when GAL4 is present only in olfactory projection neurons to drive FLP recombination (B), robust V5 staining that colocalizes with endogenous DLG is evident. (C-D) Representative confocal images of adult antennal lobes of multiple genotypes stained with antibodies to Brp (green), DLG1-V5 (red), and N-Cadherin (blue). DLG1-V5 expression is closely associated with endogenous Brp staining, though appears more apposed instead of directly colocalized, suggesting that the majority of DLG1-V5 is postsynaptic (D). (E-F) Representative confocal images of adult antennal lobes expressing Brp-Short in presynaptic DA1 ORNs and FLP-driven DLG1-V5 expression in DA1 and VA1d PNs and stained with antibodies to dsRed (green, Brp-Short-mStraw), V5 (red, DLG1-V5), and N-Cadherin (blue). *dlg1[4K]* can be combined with multiple binary expression systems in a driver-dependent manner to label multiple sets of neurons simultaneously. In this case, presynaptic active zones in DA1 ORNs are labeled via the QF system and DLG1-V5 labels postsynapses in DA1 PNs via the GAL4 system (F). In the absence of GAL4, only QF-driven expression is visible (E). Panels (A_5_, B_5_, C_5_, D_5_, E_5_) show higher magnifications of the areas within the stippled boxes (shown in A_4_, B_4_, C_4_, D_4_, E_4_) providing more detailed resolution of synaptic labeling with these components. Scale bars, 10 μm.

We next wanted to determine if we could use *dlg1[4K]* in concert with a cell-type specific presynaptic label to label the pre- and postsynaptic compartments of two different cells simultaneously. We expressed QUAS-Brp-Short using the *Or67d-QF* driver (Liang et al., 2013) to visualize presynaptic active zones in *Or67d*-positive ORNs that innervate the DA1 glomerulus. In the same brain, we labeled the PNs that are postsynaptic to *Or67d*-positive ORNs using the *Mz19-GAL4* driver (Berdnik et al., 2006) in concert with UAS-FLP and *dlg1[4K]*. In the absence of GAL4 (Fig. 4E), only QF-driven Brp-Short was evident; however, when Mz19-GAL4 supplied FLP was present, we observed DLG1 labeling only in the *Mz19*-positive PNs that innervate the DA1 and VA1d glomeruli (Fig. 4F). The *Mz19*-PN DLG1-V5 staining was closely apposed to *Or67d*-ORN Brp-Short immunoreactivity, indicating that *dlg1[4K]* successfully labeled postsynaptic regions. Further, this experiment indicated that multiple binary expression systems could be utilized in the same brain to label the pre- and postsynaptic compartments of two different cells simultaneously. These data indicate that *dlg1[4K]* is suitable for labeling adult brain postsynaptic regions with high fidelity compared to endogenous DLG1.

### Quantitative analysis of postsynaptic puncta in the antennal lobe with dlg1[4K] reveals cell-type specific patterns of synaptic organization

Distinct and stereotyped rules govern the three-dimensional synaptic organization of different classes of olfactory neurons in the antennal lobe (Mosca and Luo, 2014). These analyses, however, were largely limited to the presynaptic active zone using Brp-Short (Duhart and Mosca, 2022; Fouquet et al., 2009; Mosca and Luo, 2014; Mosca et al., 2017). Some studies have examined postsynaptic architecture using tagged acetylcholine receptors like Dα7-GFP (Christiansen et al., 2011; Kremer et al., 2010; Leiss et al., 2009; Mosca and Luo, 2014) but these analyses were limited only to one class of postsynaptic terminal and then, only a subset of that class (those containing Dα7 subunits). We used *dlg1[4K]* to assess general excitatory postsynaptic organization in different classes of antennal lobe olfactory neurons. Historically, the most well-studied connection in the antennal lobe is between ORNs and PNs. However, there is also considerable evidence of synaptic connections between ORNs and LNs as well as between LNs and PNs (Chou et al., 2010; Horne et al., 2018; Hummel et al., 2003; Rybak et al., 2016; Tobin et al., 2017). We focused on the DA1 glomerulus in the antennal lobe as there is ready genetic access to the ORNs (Liang et al., 2013; Stockinger et al., 2005), PNs (Berdnik et al., 2006; Hong et al., 2012), and multiglomerular LNs (Chou et al., 2010) that innervate DA1. We used combinations of GAL4 and QF drivers with Brp-Short (for presynaptic labeling) and *dlg1[4K]* (for postsynaptic labeling) to concurrently examine synaptic organization. Specifically, we examined three different pairs of cells representing ORN-PN (using *Or67d-QF* and *Mz19-GAL4*), ORN-LN (using *Or67d-QF* and *NP3056-GAL4*), and PN-LN synapses (using *Mz19-QF* and *NP3056-GAL4*) and quantified the number of Brp-Short and DLG1-V5 puncta in each condition and mapped the three dimensional organization of each species of puncta using nearest neighbor distance and cluster analyses (Mosca and Luo, 2014).

When we visualized ORN presynapses and PN postsynapses in DA1 (Fig. 5A-C), we observed clear apposition between presynaptic Brp-Short puncta and postsynaptic DLG1 puncta, as predicted (Fig. 5B-C) given known connectivity. We subsequently quantified DLG1 puncta in the DA1 glomerulus (Fig. 5D) and on average, DA1 PNs contain 838 ± 23 DLG1 puncta, which was slightly less than the number of Brp-Short puncta quantified within DA1 ORNs (Fig. 5D). This is consistent with connectivity patterns as multiple presynapses can be made onto a single postsynapse at ORN – PN connections in the antennal lobe (Horne et al., 2018; Mosca and Luo, 2014; Seki et al., 2010; Tobin et al., 2017). When we compared these results with other combinations of connections (ORN presynapses with LN postsynapses or PN presynapses with LN postsynapses), we found that PNs represent the predominant contribution of postsynaptic puncta within DA1, with PN postsynapses on average representing nearly double (838 ± 23 PN DLG1 puncta compared to 495 ± 28 LN DLG1 puncta for ORN-LN and 421 ± 16 LN DLG1 puncta for PN-LN) that of LN postsynapses (Fig. 5D, I, N). There was also considerably reduced apposition in both ORN-LN and PN-LN pairs when compared to ORN-PN (Fig. 5B-C, G-H, L-M), though we did observe some apposition, which is reflective of connectivity findings from EM reconstructions (Horne et al., 2018; Rybak et al., 2016). Of all three combinations, the least apposition was observed between PNs and LNs (Fig. 5L-M).

**Fig. 5.**
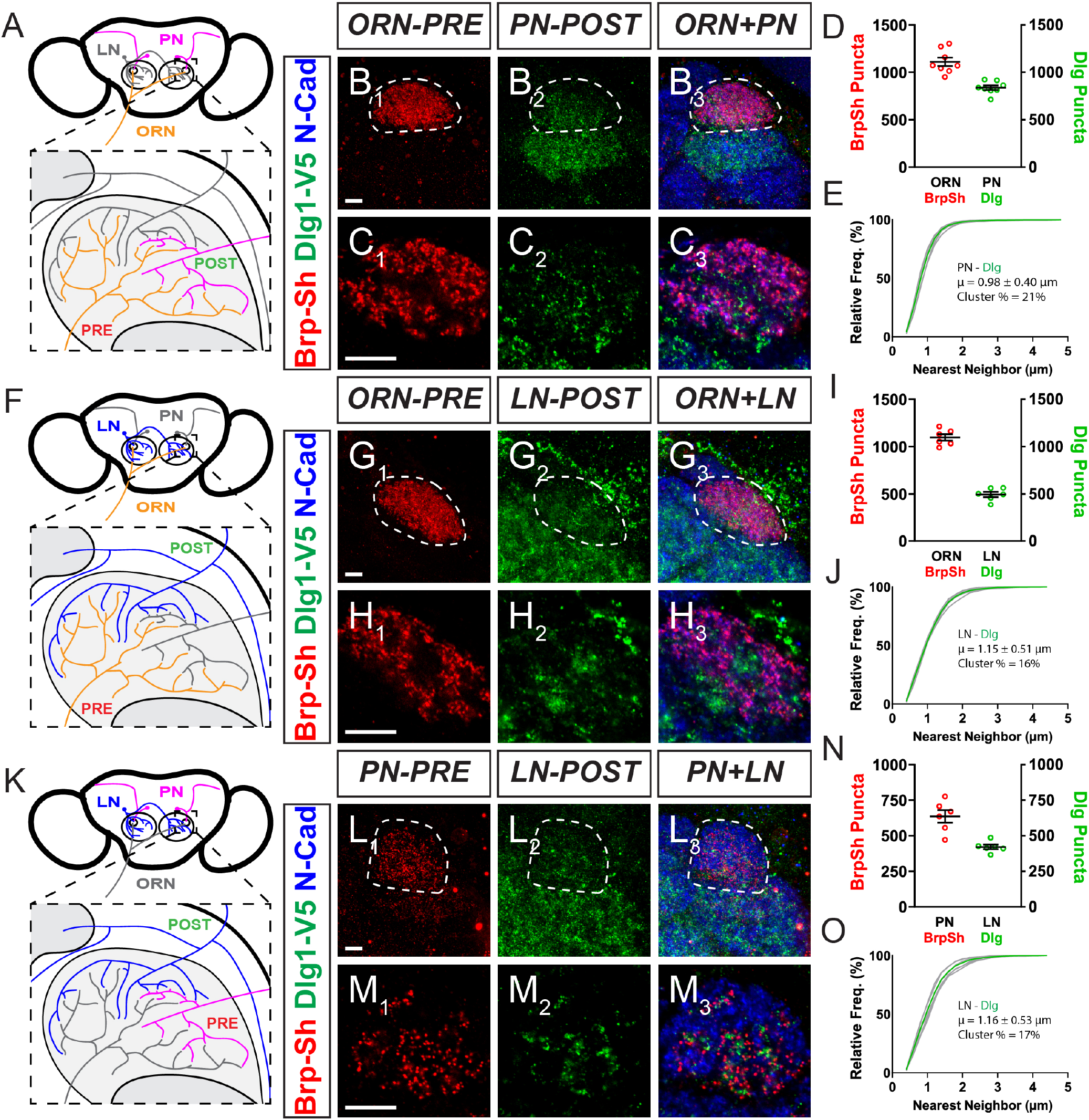
A quantitative analysis of postsynaptic DLG1 puncta for PNs and LNs in the DA1 glomerulus using *dlg1[4K]*. (A) Schematic of the *Drosophila* antennal lobes showing presynaptic ORNs (orange) and postsynaptic PNs (magenta) in the DA1 glomerulus. (B) Representative confocal maximum projections of DA1 ORNs expressing Brp-Short-mStraw and DA1 PNs expressing DLG1-V5 and stained with antibodies against mStraw (red), V5 (green), and N-Cadherin (blue). (C) Single, high magnification optical sections of the DA1 ORNs and PNs from (B). (D) Quantification of Brp-Short- mStraw puncta for ORNs and DLG1-V5 puncta for PNs. (E) Cumulative frequency histogram of the nearest neighbor distance between DLG1-V5 puncta in DA1 PNs. The average (μ) and the Cluster % of puncta with an NND between 0.6 and 0.75 μm are indicated on the graph. Gray traces represent individual glomeruli while the green trace represents the aggregate average. (F) Schematic of the *Drosophila* antennal lobes showing presynaptic ORNs (orange) and postsynaptic LNs (blue) in the DA1 glomerulus. (G-H) Representative confocal image stacks and corresponding single, high magnification sections of DA1 ORNs expressing Brp-Short-mStraw and multiglomerular LNs in the DA1 glomerulus expressing DLG1-V5, and stained with antibodies as in (B-C). (I) Quantification of Brp-Short-mStraw puncta for ORNs and DLG1-V5 puncta for LNs. (J) Cumulative frequency histogram of the nearest neighbor distance between DLG1-V5 puncta in multiglomerular LNs in DA1, including the average (μ) and the Cluster %. Traces represent individual glomeruli (gray) or the aggregate average (green). (K) Schematic of the Drosophila antennal lobes showing presynaptic PNs (magenta) and postsynaptic LNs (blue) in the DA1 glomerulus. (L-M) Representative confocal image stacks and corresponding single optical sections of DA1 PNs expressing Brp-Short-mStraw and multiglomerular LNs in the DA1 glomerulus expressing DLG1-V5 and stained with antibodies as in (B-C). (N) Quantification of Brp-Short-mStraw puncta for PNs and DLG1-V5 puncta for LNs. (O) Cumulative frequency histogram of the nearest neighbor distance between DLG1-V5 puncta in the multiglomerular LNs from (L-M), including the average (μ) and the Cluster %. Traces represent individual glomeruli (gray) or the aggregate average (green), as previously. For all conditions, n ≥ 6 glomeruli from 3 brains, and 900 (E), 675 (J), or 450 (O) individual puncta. Scale bars, 5 μm.

We also examined the three-dimensional organization of DLG1 puncta in PNs and LNs using nearest neighbor distance (NND) and clustering analyses (Mosca and Luo, 2014). Active zones in ORNs, PNs, and LNs each display distinct clustering and NND values (Mosca and Luo, 2014) that are stereotyped among brains. Interestingly, we observed a similar organization for postsynaptic DLG1 puncta (Fig. 5E, J, O). PNs displayed an NND of 0.98 ± 0.40 μm with 21% of the total puncta clustered together (Fig. 5E). This was stereotyped across multiple brains. LNs possessed an NND of 1.15 ± 0.15 μm with 16% clustered puncta (Fig. 5J) or 1.16 ± 0.53 μm with 17% clustered puncta (Fig. 5O). There was also notable stereotypy from brain to brain, similar to what we observed with PN puncta. In all, this indicates that *dlg1[4K]* can be used to quantitatively assess postsynaptic organization in multiple different classes of olfactory neurons. Moreover, we reveal that, like presynaptic active zones (Mosca and Luo, 2014), there are distinct rules that govern the three-dimensional organization of DLG1-positive postsynapses. This indicates that *dlg1[4K]* can be used similarly to Brp-Short (Duhart and Mosca, 2022) for quantitative assays of synaptic organization and development.

### dlg1[4K] can visualize a presynaptic DLG1 contribution at the NMJ

Our evidence suggests that *dlg1[4K]* has marked utility as a postsynaptic label in both the central and peripheral nervous systems, even when expressed in central neurons, which can contain both pre- and postsynaptic specializations in close proximity. However, some evidence suggests that DLG1 may also function presynaptically at peripheral NMJ synapses in *Drosophila*, and in some aspects of mammalian synaptic function (Aoki et al., 2001; Astorga et al., 2016; Budnik et al., 1996; Mendoza et al., 2003). How DLG1 functions presynaptically remains unclear; thus far only genetic evidence has suggested a role for presynaptic DLG1 in regulating the development and integrity of the subsynaptic reticulum (SSR) in muscle (Budnik et al., 1996). Presynaptic DLG1 at the NMJ has neither been successfully imaged in isolation nor separated from the predominant contribution of postsynaptic DLG1.

To determine if *dlg1[4K]* could be used to specifically visualize presynaptic DLG1 at peripheral synapses, we expressed FLP recombinase in all neurons (via *C155-GAL4*; Lin and Goodman, 1994) in *dlg1[4K]* larvae and imaged NMJ boutons. In the absence of GAL4, we observed no DLG1-V5 staining (Fig. 6A). However, when GAL4 was present to catalyze FLP recombination, we observed robust DLG1-V5 staining restricted to presynaptic boutons (Fig. 6B). DLG1-V5 immunoreactivity was concentrated in and filled synaptic boutons, overlapping with HRP immunoreactivity, which recognizes insect neurons (Jan and Jan, 1982). High magnification imaging of boutons showed that DLG1-V5 staining resembled synaptic vesicle staining in boutons and was largely (but not completely) excluded from the inter-bouton axon (Fig. 6C). When DLG1-V5 was examined in larval lysates via Western blot, a neuronal contribution could be visualized with extended exposure of the blot (Fig. 6D). Under such conditions, the signal produced by the muscle was vastly oversaturated, indicating that the predominant pool of DLG1 at the NMJ is postsynaptic. This is consistent with DLG1 immunostaining in presynaptic *dlg1[4K]* larvae, which partially overlapped with the presynaptic bouton (Fig. 6E) but was largely postsynaptic. There was no significant overlap with α-spectrin (Fig. 6F), which largely labels the postsynaptic region of NMJ boutons (Pielage et al., 2006). Taken together, these results, for the first time, visualize presynaptic DLG1 *in vivo* at the NMJ, supporting genetic evidence for a functional role. Moreover, our results demonstrate, as suggested by endogenous DLG1 staining that the predominant contribution of DLG1 is postsynaptic, but presynaptic DLG1 may be involved throughout the bouton, potentially with synaptic vesicles.

**Fig. 6.**
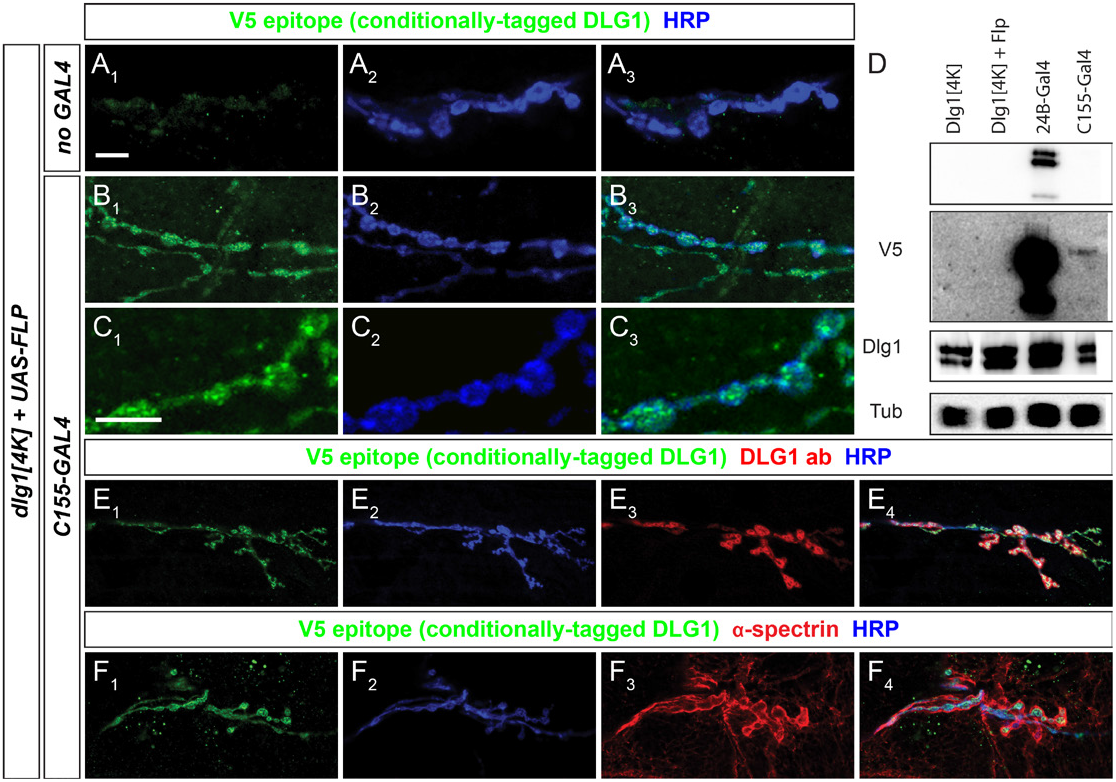
Presynaptic DLG1 in motoneurons can be isolated and visualized using *dlg1[4K]*. (A-C) Representative confocal images of larvae with the *dlg1[4K]* allele and FLP recombinase lacking (A) or with (B-C) a pan-neuronal GAL4 source and stained for antibodies to DLG1-V5 (green) and HRP (blue). In the absence of GAL4 (A), no labeling is observed, but when GAL4 is present, presynaptic DLG1-V5 (B-C) is evident in presynaptic motoneuron terminals at lower (B) and higher (C) magnification. (D) Western blot analysis of multiple genotypes activating *dlg1[4K]* in muscle or neurons. In an absence of a GAL4-driven FLP source, no V5 is evident. When FLP is provided in muscle or nerve, at short exposures, V5 is only evident in muscle (top) but at long exposures (2nd panel), neuronal DLG1-V5 can be observed. Tubulin and endogenous DLG1 are used as loading controls. (E-F) Representative confocal images of larvae with DLG1-V5 labeling in motoneurons and stained with antibodies to DLG1-V5 (green), endogenous DLG1 (red, E), α-spectrin (red, F), and HRP. Some overlap can be seen (E) with endogenous DLG1 staining while there is clear distinction at boutons between DLG1-V5 and postsynaptic α-spectrin staining (F). Scale bars, 10 μm.

### dlg1[4K] enables visualization of extra-neuronal DLG1

Beyond the nervous system, DLG1 in *Drosophila* also functions in tumor suppression, formation of septate junctions, oocyte biology, and epithelial cell polarity (Bilder et al., 2000, 2003; Su et al., 2013; Tepass et al., 2001; Woods and Bryant, 1991; Woods et al., 1996). We reasoned that *dlg1[4K]* would also be advantageous in visualizing cell-type specific contributions of DLG1 to those processes as well. To examine this, we tested whether *dlg1[4K]* could reveal DLG1 expression in non-neuronal tissues, specifically, the ovarian follicle epithelia, a tissue where DLG1 has a stereotyped, distinctive expression pattern and contributes to establishing planar cell polarity (Bilder et al., 2000). In the ovary, DLG1 recognizes the follicle cells that line the perimeter of the oocyte and the borders of the nurse cells (Fig. S4A). The GR1-GAL4 driver has been used to investigate the role of dlg1 in the ovary and specifically expresses in the follicle cells but not the nurse cells or oocyte (Tran and Berg, 2003). When we drove FLP expression using *GR1-GAL4* in the *dlg1[4K]* background, we observed robust V5 signal only in the follicle cells that precisely colocalized with endogenous DLG1 staining (Fig. S4B). This demonstrates that *dlg1[4K]* can also be used in non-neuronal tissues to visualize endogenous DLG1 expression. In all, *dlg1[4K]* is a powerful tool for examining both quantitative and qualitative expression of endogenous DLG1 throughout the fly with cell-type specificity.

## DISCUSSION

The organization of synaptic connections in neural circuits is a key determinant of how those circuits function, drive behavior, and enable communication from one cell to another. The three-dimensional organization of synapses underlies neural computation and is essential for normal function (Montero-Crespo et al., 2020; Turner et al., 2022; Xu et al., 2021; Zhan et al., 2016). Moreover, understanding how synapses in circuits are organized offers a window into understanding the paradigms that govern synaptic development and a foundation for how a nervous system is assembled. This blueprint for development informs how synaptic organization is disrupted by neurodevelopmental, neuropsychiatric, and even neurodegenerative diseases. In *Drosophila*, a number of techniques are aimed at studying three-dimensional synapse organization (Chen et al., 2014; Duhart and Mosca, 2022; Landínez-Macías et al., 2021; Mosca and Luo, 2014; Mosca et al., 2017; Urwyler et al., 2015, 2019) but largely focus on general presynaptic active zone markers. A necessity for the thorough study of synaptogenesis involves distinguishing the pre- vs postsynaptic elements of the synapse in a directed subset of neurons to examine connectivity and assess experimental outcomes. The diversity of postsynaptic structures has made a generalized marker difficult to develop, particularly one that functions in a specific set of target neurons. Some strategies used tagged neurotransmitter receptors (Andlauer et al., 2014; Chen et al., 2014; Kremer et al., 2010; Leiss et al., 2009; Mosca and Luo, 2014) to examine synaptic organization but those strategies often relied on overexpression and / or only examine a subset of postsynapses. Newer strategies use conditionally modified endogenous versions (Fendl et al., 2020) to label neurotransmitter receptors with cell-type specificity under the expression control of their endogenous promoters to circumvent issues of overexpression but still only label select postsynapses.

To produce a general excitatory postsynaptic label that was cell-type specific and under the control of its endogenous promoter, we designed a conditional strategy based on FLP recombination and used CRISPR/Cas9 genome editing to modify the dlg1 locus in *Drosophila*. dlg1 encodes a homologue of the well-known mammalian postsynaptic protein PSD-95 (Won et al., 2017) and is an established postsynaptic protein in *Drosophila* (Budnik et al., 1996; Gorczyca et al., 2007; Rivlin et al., 2004). By inserting an FRT-STOP-FRT-V5 tag immediately before the endogenous STOP codon of the dlg1 gene (Fig. 1), we created *dlg1[4K]*, which enables endogenously expressed DLG1 to be tagged with a V5 epitope only in tissues where dlg1 is endogenously expressed and FLP is present to excise the preceding STOP cassette. As proof-of-principle, we demonstrated that *dlg1[4K]* specifically labeled DLG1 with multiple binary expression systems and at multiple peripheral and central synapses where *dlg1* is known to be expressed (Figs. 2–4). Cell-type specific experiments thus validated the utility and versatility of *dlg1[4K]* in glutamatergic and cholinergic neurons, further highlighting the generality of this postsynaptic marker. When DLG1 is isolated in neurons, it can specifically label postsynapses (Fig. 2, 5) and we find that distinct rules govern the quantitative and qualitative three-dimensional organization of postsynaptic terminals (Fig. 5). Finally, we visualize, for the first time, DLG1 presynaptic expression at the NMJ, where previous genetic evidence had intimated a role in synaptic organization. In all, we present a new conditional strategy to generally label postsynapses with cell-type specificity that we anticipate will be broadly useful to the study of synaptic organization.

We used *dlg1[4K]* to more closely examine postsynaptic organization in the neurons that comprise the antennal lobe, the first order processing center of olfactory information in the *Drosophila* brain. The fly antennal lobe is a powerful system for examining synaptic organization (Duhart and Mosca, 2022; Mosca and Luo, 2014; Mosca et al., 2017). Presynaptic organization in antennal lobe neurons follow distinct rules that govern active zone clustering, distance, and density (Mosca and Luo, 2014) that are stereotyped depending on neuron class. Our understanding of postsynaptic organization, however, has been limited. Previous work examined acetylcholine receptor organization (Mosca and Luo, 2014) but this is incomplete as it only examines one potential class of postsynaptic receptors. *dlg1[4K]* enabled us to examine the general postsynaptic organization in PNs and LNs and, in combination with multiple binary expression systems, assess the concurrent organization of presynaptic active zones from ORNs or PNs (Fig. 5). We discovered differences in both qualitative and quantitative aspects of postsynaptic organization in different classes of olfactory neurons, similar to our previous work on presynaptic organization (Mosca and Luo, 2014). PN and LN postsynapses displayed distinct subglomerular organization whether considered independently or with respect to different types of labeled (ORN or PN) presynapses (Fig. 5). Further, PN and LN postsynapses displayed stereotyped nearest neighbor distances as well as clustering percentages that differed from each other (Fig. 5). This indicates that, as for presynaptic active zones, olfactory neuron classes use distinct rules to organize postsynapses in three-dimensions. The stereotypy suggests that some aspects of these rules may be hardwired; future genetic studies will be very informative to assess how organization is accomplished in the CNS. Overall, this indicates that *dlg1[4K]* is useful for the quantitative and three-dimensional analysis of CNS postsynaptic organization.

At NMJ synapses, DLG1 plays a predominantly postsynaptic role in synapse maturation, organization, and in building ultrastructure like the SSR (Chou et al., 2020; Gramates and Budnik, 1999; Harris and Littleton, 2015). However, genetic evidence suggests that a presynaptic pool of DLG1 (Astorga et al., 2016; Budnik et al., 1996) regulates neuronal function. To date, presynaptic DLG1 has not been observed in isolation, largely because any potential signal is likely occluded by the predominant postsynaptic immunohistochemical staining of DLG1 at the NMJ. Using *dlg1[4K]*, we isolated the presynaptic pool of DLG1 and showed that it localizes throughout the presynaptic bouton (Fig. 6). The significance of this data is twofold. First, it offers the first visualization of presynaptic DLG1 at the NMJ, enabling the study of that pool in isolation including subsynaptic localization. Second, it raises an important potential caveat about *dlg1[4K]*. In data from the NMJ (Figs. 2, 6), the larval ventral nerve cord (Fig. 3), olfactory neurons in the CNS (Figs. 4, 5), and from the ovary (Fig. 4), *dlg1[4K]* accurately reflects the endogenous expression pattern of DLG1. Predominantly, that localization is postsynaptic, but as the data from the NMJ shows, it can be presynaptic. Therefore, care must be taken when interpreting data from *dlg1[4K]*. Though the localization of *dlg1[4K]* in the CNS is consistent with a predominantly, if not completely, postsynaptic localization (Fig. 3–5), we cannot rule out a potential presynaptic contribution. This is reflective of DLG1 biology; therefore, careful co-localization and genetic evidence should be used in concert with *dlg1[4K]* to rule out any potential presynaptic contributions. Our evidence suggests that *dlg1[4K]* has great utility in the CNS as a postsynaptic marker but is ultimately reflective of the endogenous expression of *dlg1*, wherever that may be.

DLG1 functions in many diverse tissues including and beyond the nervous system and plays multiple roles in cell adhesion, synaptic organization, and cell polarity (Bilder et al., 2000, 2003; Khoury and Bilder, 2022; Su et al., 2013; Tepass et al., 2001; Woods and Bryant, 1991). The broad functional roles of DLG1 are mediated by several translational isoforms generated from alternative STOP codons (Fig. S1). As the *dlg1[4K]* strategy integrates an FRT-STOP-FRT-V5 at the distal most STOP codon (Fig. 1), this raises the important caveat that 7 isoforms of DLG1 will not be labeled by this strategy. *dlg1[4K]* labels isoforms that include protein motifs known to function at the postsynapse (Won et al., 2017) as our goal was to create a cell-type specific conditionally expressed postsynaptic label. The L27, three PDZ, SH3, and GK domains that most closely resemble vertebrate DLG1-4 orthologs (Won et al., 2017) are included; some of the 7 isoforms not labeled exclude these protein motifs. Specifically, the L27 domain at the N-terminus of DLG1 is thought to be important in forming supramolecular complexes that may allow homomultimerization of DLG1/SAP97 in mammalian postsynapses to influence neurotransmitter receptor organization (Nakagawa et al., 2004). Ten *Drosophila* isoforms contain the L27 domain but 5 of those 10 exclude the other domains (Graveley et al., 2011) that are required for postsynaptic function. Therefore, those 5 isoforms will not be labeled by *dlg1[4K]* but the remaining L27-containing isoforms will be labeled. The isoform structure of DLG1 in other tissues also remains incompletely understood so it remains an important caveat of *dlg1[4K]* to be cognizant of the important isoforms used in the process being studied. For neuronal purposes, our evidence suggests that isoforms relevant to postsynaptic function are labeled successfully by *dlg1[4K]*, supporting its use.

Understanding postsynaptic organization at the level of individual neuron types has lagged behind the study of presynaptic organization due to a lack of suitable general tools. Work with tagged neurotransmitter receptors has been invaluable for understanding synaptic organization (Andlauer et al., 2014; Chen et al., 2014; Fendl et al., 2020; Leiss et al., 2009; Mosca and Luo, 2014) but is necessarily limited to a single class of (mostly excitatory) synapses. *dlg1[4K]* represents the first general postsynaptic label in *Drosophila* that encompasses multiple types of postsynapses. We anticipate that this strategy will be widely usable to understand postsynaptic organization in many different classes of neurons, without the negative caveats that can be associated with overexpression or impenetrable cell density. Moreover, we also envision the strategy as expandable. Though *dlg1[4K]* includes an epitope tag, this can be modified to include a fluorescent reporter, any kind of general effector, or even a degron (Okoye et al., 2022) to study the targeted function of DLG1. Moreover, by adjusting the homology arms of the construct (Fig. 1), this strategy can be applied to tagging most any gene with amenable PAM sites at the C-terminus. Further, the strategy could be generally applied like current FLP technologies (Fendl et al., 2020) including FLPStop (Fisher et al., 2017) to modify endogenous genes within the coding sequence. Such aspects will notably advance studying synaptic organization using both qualitative and quantitative assays and allow for a deeper understanding of three-dimensional postsynaptic organization. By first understanding the foundation of postsynaptic biology, we can better grasp how it is influenced by neurodevelopmental and neurodegenerative disease and better probe underlying disease mechanisms.

## ACKNOWLEDGMENTS

We’d like to thank all members of the Mosca Lab for continued support, discussion, and critical assessment of the project throughout its undertaking. We would like to thank Dr. Kristen Davis, Alison DePew, Dr. Juan Duhart, Jesse Humenik, and S. Zosimus for comments on the manuscript. We are also grateful to the fly community for making so many integral resources available, including FlyBase and FlyCRISPR. Stocks obtained from the Bloomington *Drosophila* Stock Center (NIH P40OD018537) were used in this study. We appreciate the availability of monoclonal antibodies used in this study from the Developmental Studies Hybridoma Bank, (created by the NICHD of the NIH and maintained at The University of Iowa, Department of Biology, Iowa City, IA 52242). This work was supported by US National Institute of Health grants R01-NS110907 and R00-DC013059 (to TJM), Commonwealth Universal Research Enhancement of the Pennsylvania Department of Health grant 4100077067 (to TJM). Work in the TJM lab is supported by the Alfred P. Sloan Foundation, the Whitehall Foundation, the Jefferson Synaptic Biology Center, and TJU start-up funds.

## AUTHOR CONTRIBUTIONS

M.J.P. and T.J.M. designed the project; M.J.P. and M.A. performed experiments; M.P. and T.J.M. produced reagents; M.J.P. and M.A. analyzed the data. M.J.P., M.A., and T.J.M. wrote and edited the manuscript.

## MATERIALS AND METHODS

### Lead Contact

Requests for any resources or reagents should be addressed to the Lead Contact, Timothy J. Mosca (timothy.mosca@ jefferson.edu).

### Materials Availability

All plasmids, transgenic flies, antibodies, and custom reagents created for this study are available upon request to the Lead Contact.

### Data Availability

Original data including image files, Western blots, or statistical analyses are available from the Lead Contact on reasonable request.

### Construction of the dlg1[4K] Plasmid and Transgenic Line

To introduce a conditionally expressed tag with recombinase sites into *dlg1*, we inserted a 3.34 kB construct directly into the endogenous *dlg1* locus using homology directed repair (HDR) of a donor plasmid and CRISPR/Cas9 genome editing (Gratz et al., 2013, 2015; Port et al., 2014) at a PAM site 12 bp upstream of the most distal stop codon. We constructed the *dlg1[4K]* donor plasmid (Fig. 1) by joining five fragments into a minimal (AMP and ORI) plasmid backbone by In Fusion assembly (Takara, no. 639649). A list of primers used, and precise descriptions of the fragments, are found in Table 2. Each fragment was amplified using Q5 polymerase (New England Biolabs, no. M0491s) with custom primers (IDT, Coralville IA). The fragments included the AMP ORI backbone and an *FRT>STOP>FRT* sequence, both derived from *pJFRC203-10XUAS-FRT>STOP>FRT-myr∷smGFP-cMyc*. (RRID: Addgene 63167). We also added a DsRed visible fluorescent marker cassette, derived from *pScarlessHD-2xHA-DsRed* (which was a gift from Kate O’Connor-Giles; Addgene plasmid #80822; RRID:Addgene_80822) to mark any resulting transformants. We inserted the DsRed cassette, which itself is flanked by PiggyBac inverted repeats, at a TTAA sequence within the *FRT>STOP>FRT* cassette. This positioning enabled us to use the DsRed to identify transformants but then remove later by crossing transformant lines to a PiggyBac transposase source. The DsRed cassette would be removed in a scarless fashion, leaving no additional sequence (as it was cloned directly into a homologous matching TTAA sequence in the *FRT>STOP>FRT* cassette. An additional fragment including the 3’ end of the *FRT>STOP>FRT* cassette, a 3x- V5 epitope tag, and homologous sequence that extended into the ‘scarless’ cassette was synthetically manufactured (GenScript). Finally, 500 bp left and right homology arms flanking the double strand break that would be induced at the PAM site were amplified directly from the *Vasa-Cas9* injection strain (BL51324) to ensure sequence compatibility. The left (5’) homology arm was placed directly in front of the 5’-most FRT site and we added a single C nucleotide inserted directly upstream of the FRT sequence to maintain an open reading frame after the FLP-out recombination event. The right homology arm was positioned exactly at the Cas9 cut site; this left 12 bp of the *dlg1* coding sequence between the 3’ most FRT sequence and the V5 epitope tag, which encodes 4 amino acids of the DLG1 carboxy terminus. Intermediate cloning steps and the final donor plasmid were sequence verified (GeneWiz, South Plainfield NJ). Donor construct sequence is available upon request.

To perform CRISPR/Cas9 at the *dlg1* locus, *vasa-Cas9* embryos (BDSC 51324) were injected (BestGene, Chino Hills CA) with the donor plasmid and a guide RNA (gRNA) plasmid; the gRNA was made by annealing sense and antisense oligos homologous to sequence adjacent to and 5’ of the targeted *dlg1* PAM site (Table 2) and cloned into *pU6-BbsI-chiRNA* (Gratz et al., 2013). Transformant lines expressing 3xP3 DsRed in the eye were identified visually and then verified for correct integration into *dlg1* by amplifying and sequencing genomic DNA spanning the homology arm breakpoints. This verified that the knocked-in sequence was at the predicted position with no genomic rearrangements or nucleotide substitutions. Transgenic knock-in lines received from BestGene were crossed to *Herm{3xP3-ECFP, α-tub-piggyBacK10}M6* (BL55804) to excise the DsRed cassette. DsRed negative progeny were balanced and sequence verified to demonstrate precise excision of the scarless cassette. A single line (*dlg1[4K]*) was chosen and homozygous females used for further experimentation.

### Drosophila stocks and transgenic lines

All *Drosophila* stocks and crosses were grown on cornmeal medium (Archon Scientific, Durham, NC) at 25°C and 60% humidity with a 12/12 light/dark cycle in specialized incubators (Darwin Chambers, St. Louis, MO). All alleles, GAL4 drivers, and UAS lines were maintained over phenotypically selectable balancer lines to ensure facile identification. The *dlg1[4K]* line was established over an FM7 balancer chromosome and subsequently utilized in all experiments and recombinations. The following GAL4, QF, or LexA lines were used to enable tissue-specific expression: *DMef2-GAL4* (Lilly et al., 1995) (pan-muscle expression), *elav^C155^-GAL4* (Lin and Goodman, 1994) (pan-neuronal expression), *how24B-GAL4* (Brand and Perrimon, 1993) (pan-muscle expression), *BG487-GAL4* (Budnik et al., 1996) (muscle subset expression), *GH146-GAL4* (Stocker et al., 1997) (olfactory projection neuron subset expression), *Mz19-GAL4* (Jefferis et al., 2004) (DA1, VA1d, and DC3 olfactory projection neuron expression), *NP3056-GAL4* (Chou et al., 2010) (multiglomerular local interneuron expression), *Or67d-GAL4* (Kurtovic et al., 2007) (DA1 olfactory receptor neuron expression), *GR1-Gal4* (Tran and Berg, 2003) (ovary expression), *GH146-LexA* (Lai and Lee, 2006) (olfactory projection neuron subset expression), *DMef2-LexA* (Pfeiffer et al., 2010) (pan-muscle expression), *DMef2-QF2* (Lin and Potter, 2016) (pan-muscle expression), *Or67d-QF* (Liang et al., 2013) (DA1 ORN expression), and *Synaptojanin-QF* (Petersen and Stowers, 2011) (pan-neuronal expression). The following UAS transgenes were used: *UAS-FLP* (Thibault et al., 2004), *UAS-FLP* (Nern et al., 2011), *lexA-op-FLP* (Pfeiffer et al., 2010), *QUAS-FLP* (Potter et al., 2010), *UAS-Brp-Short-mStraw* (Fouquet et al., 2009), *QUAS-Brp-Short-mStraw* (Mosca and Luo, 2014). In all experiments, homozygous *dlg1[4K]*, *UAS-FLP* recombinant females were crossed to GAL4, QF, or LexA drivers or outcrossed to *w[1118]* (BL5905) as ‘no driver’ controls. Specific genotypes are indicated in Table 1.

### NMJ, Brain, and Ovarian Tissue Immunohistochemistry

Larvae were processed for antibody staining as described (Mosca and Schwarz, 2010; Restrepo et al., 2022). Wandering third instar larvae were grown in population cages (Genesee, no. 59-100) on grape juice plates supplemented with yeast paste and then dissected in Ca^2+^-free modified *Drosophila* saline (White et al., 2001). Where driver lines were X-linked, we selected only female larvae for experimentation to ensure the presence of all genetic components. For adult flies, brain dissections were done according to (Wu and Luo, 2006) and dissected in PBST (phosphate buffered saline with 0.3% Triton-X-100) and the tracheae removed. Ovaries were dissected from three-day old adult females in PBST.

All samples (larval, brain and ovary) were fixed in 4% paraformaldehyde in 1X PBST for 20 minutes followed by three 20-minute washes in PBST. Adult samples were blocked in 5% normal goat serum and incubated with primary and secondary antibodies for two days each at 4°C. Larval samples were incubated in primary antibodies overnight at 4°C and in secondary antibodies on the subsequent day for 2h at RT. The following primary antibodies were used: mouse anti-Dlg (DSHB, cat. no. mAb4F3, 1:500) (Parnas et al., 2001), mouse anti-α-spectrin (DSHB, cat. no. mAb3A9, 1:50) (Byers et al., 1987), mouse anti-Brp (DSHB, cat. no. mAbnc82, 1:250) (Laissue et al., 1999), mouse anti-CSP (DSHB, cat. no. mAb6D6, 1:100) (Zinsmaier et al., 1994), rabbit anti-dsRed (TaKaRa Bio, cat. no. 632496, 1:250), chicken anti-GFP (Aves, cat.no. GFP-1020, 1:1000), rat anti-N-Cadherin (DSHB, cat. no. mAbDNEX-8, 1:40) (Iwai et al., 1997), mouse anti-V5 (Sigma, cat. no. SAB2702199, 1:100), mouse anti-V5 (ThermoFisher, cat. no. MA515253, 1:100), rabbit anti-V5 (ThermoFisher, cat. no. PA1993, 1:100), rabbit anti-V5 (Cell Signaling, cat. no. 13202S, 1:100), Alexa647-conjugated goat anti-HRP (Jackson ImmunoResearch, cat. no. 123-605-021, 1:100). Alexa488 and Alexa647-conjugated (Jackson ImmunoResearch, cat. nos. 715-545-151, 712-545-153, 711-545-152, 712-605-153), and Alexa 568-conjugated (ThermoFisher, cat. nos. A-11004, A-11011, A-11077, A-11041) secondary antibodies were used at 1:250. FITC-conjugated (Jackson ImmunoResearch, cat. no.703-095-155) was used at 1:200. Alexa647-conjugated phalloidin (ThermoFisher, cat.no. A22287) was used at 1:300. Samples processed for imaging were mounted in Vectashield (larval NMJs) or Slowfade (brains and ovaries) and imaged on a Zeiss LSM880 Laser Scanning Confocal Microscope (Carl Zeiss, Oberlochen, Germany). Images were further processed and figures constructed using ZEN 2.3 software (Carl Zeiss, Oberlochen, Germany), Adobe Photoshop 2022, and Adobe Illustrator 2022 (Adobe Systems, San Jose, CA).

### Imaging and Quantification of DLG1-V5 Puncta in the Brain

All images of olfactory glomeruli were obtained using a 40X 1.4 NA Plan-Apochromat lens or a 63X 1.4 NA Plan-Apochromat f/ELYRA lens at an optical zoom of 3x. Images were centered on the glomerulus of interest and the z-boundaries were set based on the appearance of the two synaptic labels - Brp-Short-mStraw and DLG1-V5. Images were analyzed three dimensionally using the Imaris Software 9.7.1 (Oxford Instruments, Abingdon, UK) on a custom-built image processing computer (Digital Storm, Fremont, CA) following previously established methods (Mosca and Luo, 2014; Mosca et al., 2017; Restrepo et al., 2022). Both Brp-Short and DLG1-V5 puncta were quantified using the “Spots” function with a spot size of 0.6 μm. The resultant masks were then visually inspected to ensure their conformation to immunostaining.

To calculate nearest neighbor distance (NND), we used “Object-Object Statistics” as part of the “Spots” function for both Brp-Short puncta and DLG1[4K] puncta. The individual values for “DistMin” and cumulative frequency histograms were obtained from Imaris and compiled in Prism 8 (GraphPad Software, Inc., La Jolla, CA). Because each punctum is about 0.6 μm in diameter, the minimum NND possible for two immediately adjacent puncta should be 0.6. Therefore, we defined clustering as puncta with an NND between the minimum possible value (0.6 μm) and 1.25 x the minimum possible NND (0.75 μm). The “Cluster%” was calculated by dividing the number of puncta with an NND value between 0.6 and 0.75 by the total number of puncta. Images were processed and figures produced using ImageJ (NIH, Bethesda, MD), Adobe Photoshop 2022, and Adobe Illustrator 2022 (Adobe Systems, San Jose, CA).

### Western Blot Analysis

For all protein samples, 20 third instar larvae per genotype were flash frozen in liquid nitrogen and homogenized in 100 μL RIPA buffer (Cell Signaling) supplemented with cOmplete protease inhibitor cocktail (Roche, no. 11873580001). Following homogenization, an additional 300 μL of RIPA buffer were added and samples were centrifuged for 10 minutes at 17,900 x g at 4°C. The resultant supernatant was removed and subsequently diluted in equal volumes of 2X SDS loading buffer (100 mM Tris pH 6.0, 4% w/v SDS, 0.2% (w/v) bromophenol blue, 5% β-mercaptoethanol) to be run on SDS-PAGE gels using the Mini Protean system (Bio-Rad, no. 1658004). Samples were heated at 95°C for 5 minutes and loaded on 4-15% TGX gels (Bio-Rad, no. 4568083) in running buffer (25 mM Tris, 192 mM glycine, 0.1% SDS) and run at 100V until complete. The resultant gels were transferred to nitrocellulose (Bio-Rad, no. 1620112) in transfer buffer (25 mM Tris, 192 mM glycine, 20% methanol) for 1 hour at 350 mA. Blots were blocked with 5% nonfat dry milk in 1X PBS and incubated with primary antibodies overnight at 4oC. The following primary antibodies were used: mouse anti-V5 (ThermoFisher, cat. no. MA51523 1:1000), mouse anti-Tubulin (Sigma, cat. no. T6199, 1:1000) and mouse anti-DLG1 (DSHB cat. no. mAb4F3, 1:1000) (Parnas et al., 2001). All blots were processed for chemiluminescence by incubation with HRP-conjugated donkey anti-mouse secondary antibodies (Jackson ImmunoResearch cat. no. 715-035-151, 1:75,000) for 2 hours at 22°C. Western blots were developed with SuperSignal West Femto substrate (ThermoFisher, no. 34095) and imaged on an Azure 400 imager (Azure Biosystems).

---

**Fig. S1.**
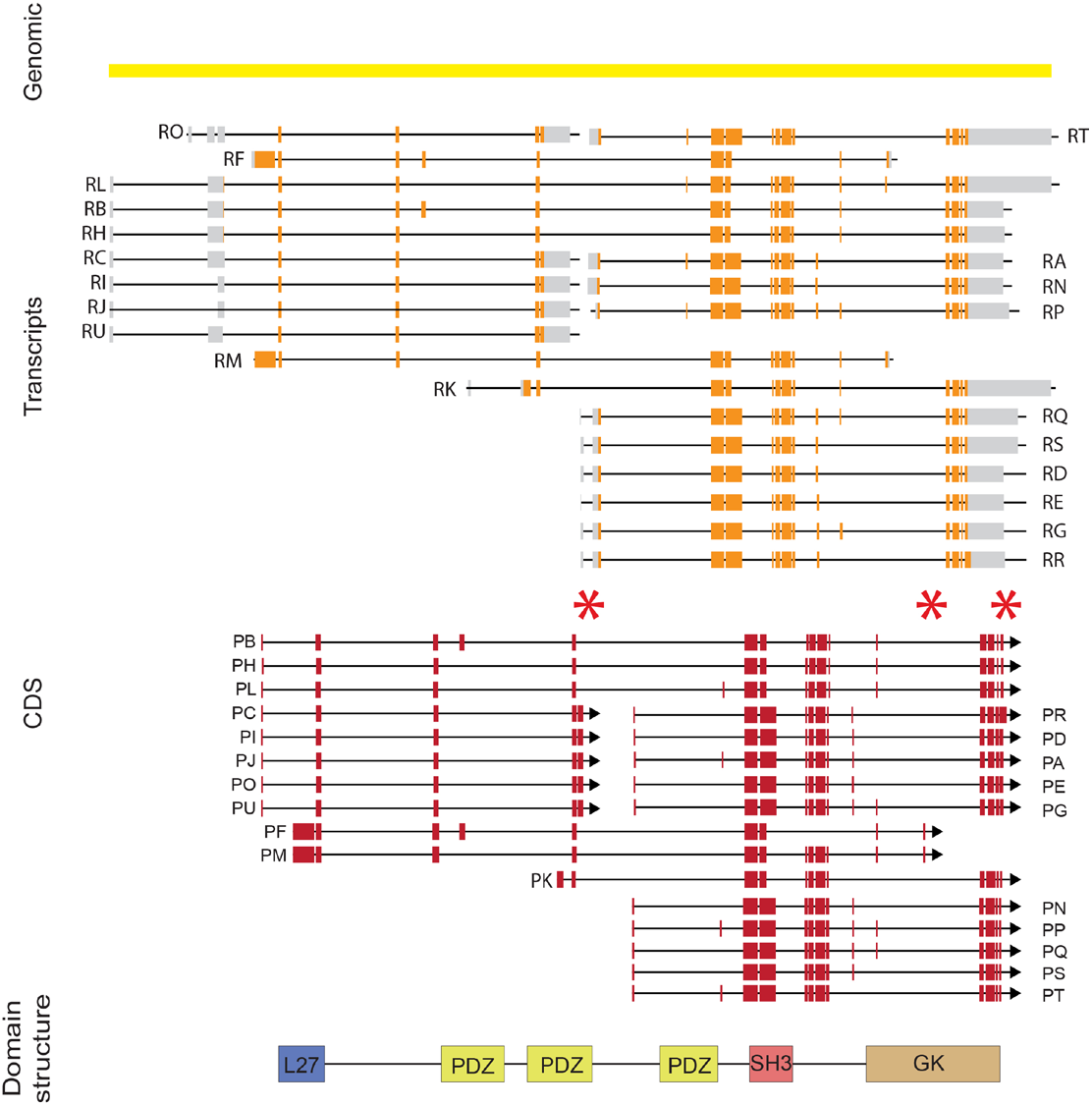
Isoform complexity at the *dlg1* locus. *dlg1* transcripts and CDS isoforms (adapted from FlyBase using version FB2022_04, released 8 August 2022). Asterisks indicate three alternative stop codons utilized by different protein isoforms. Bottom is a schematic showing protein interaction domains.

**Fig. S2.**
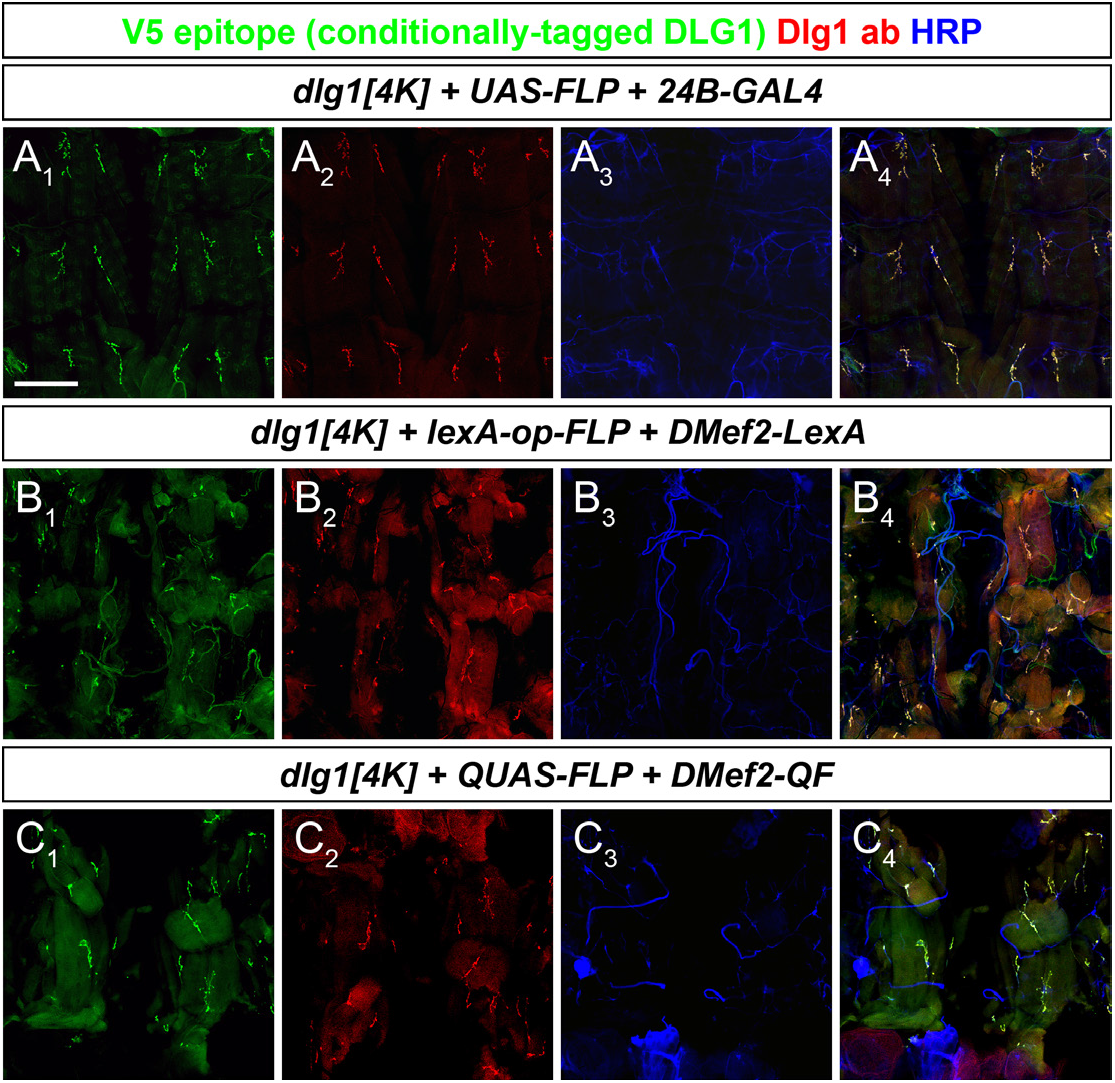
Additional examples of *dlg1[4K]* expression in larval muscles and muscle subsets. (A-C) Representative confocal images of multiple genotypes expressing FLP via the GAL4 (A), LexA (B), and QF (C) system to catalyze *dlg1[4K]* recombination and labeling and stained for antibodies to DLG1-V5 (green), endogenous DLG1 (red), and HRP (blue). In each case, staining of DLG1-V5 precisely overlaps with endogenous DLG1 throughout the larval musculature. Scale bar, 200 μm

**Fig. S3.**
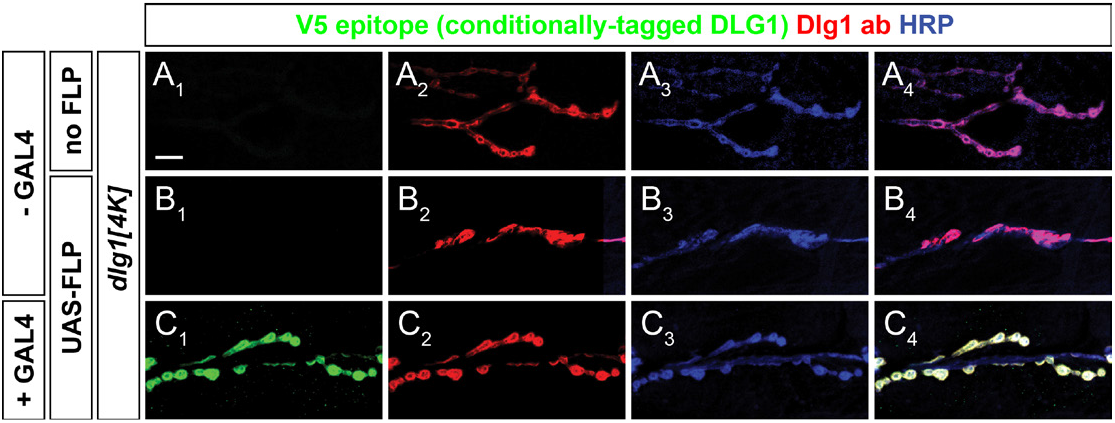
*dlg1[4K]* shows tight coupling of DLG1-V5 expression to the presence of GAL4-driven FLP. Representative confocal images of NMJs in multiple genotypes and stained with antibodies to DLG1-V5 (green), endogenous DLG1 (red), and HRP. In the absence of both GAL4 and FLP (A) or only FLP but no GAL4 (B), only endogenous DLG1 is evident. When GAL4 and UAS-FLP are present (C), DLG1-V5 expression is robustly observed. This indicates that there is little “leak” expression associated with *dlg1[4K]* V5 labeling. Scale bar, 10 μm.

**Fig. S4.**
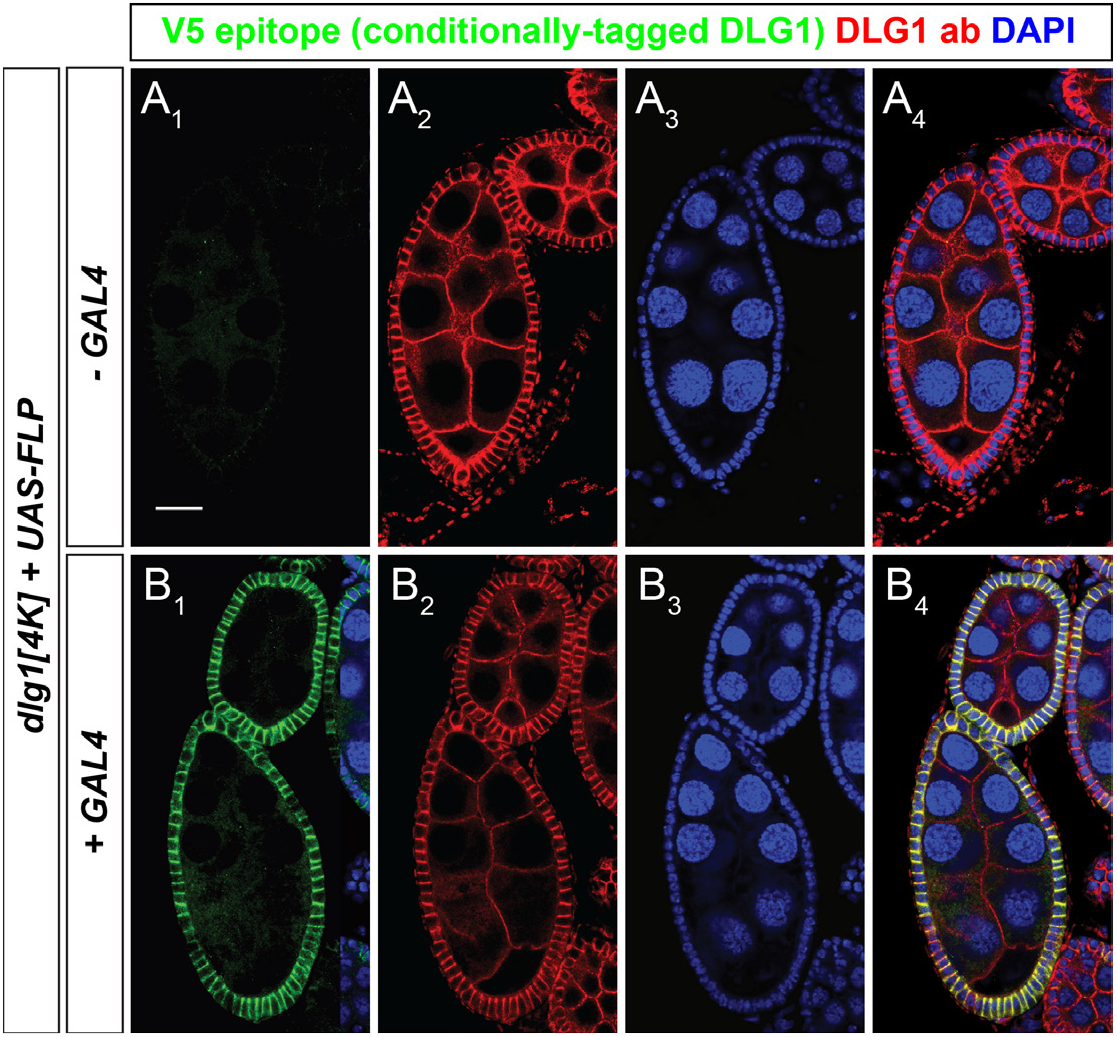
Expression of DLG1-V5 via *dlg1[4K]* in a non-neuronal cell type. (A-B) Representative confocal images of three-day old, early to mid-stage egg chambers of *dlg1[4k]* flies with a *UAS-FLP* transgene in either the absence (A) or presence (B) of GR1-GAL4 in the ovarian follicular epithelia and stained with DAPI (blue) and antibodies to DLG1-V5 (green) and endogenous DLG1 (red). In the absence of GAL4 (A), no DLG1-V5 is evident but when GAL4 is present (B) DLG1-V5 expression is seen in the follicular epithelia and precisely overlaps with endogenous DLG1 staining. Note that in these experiments, GAL4 is not expressed in the germline so DLG1-V5 signal is not observed with endogenous DLG1 in oocyte and nurse cell membranes. Scale bar, 20 μm.

